# Wounding activates the HSFA1 transcription factors to promote cellular reprogramming in Arabidopsis

**DOI:** 10.1101/2025.09.29.679115

**Authors:** Duncan Coleman, Akira Iwase, Ayako Kawamura, Arika Takebayashi, Katja E. Jager, Maolin Peng, Yutaka Kodama, David S. Favero, Tatsuya Takahashi, Momoko Ikeuchi, Takamasa Suzuki, Naohiko Ohama, Kazuko Yamaguchi-Shinozaki, Philip A. Wigge, Lieven de Veylder, Keiko Sugimoto

## Abstract

Mechanical injury is a primary trigger for cellular reprogramming during organ regeneration, yet the molecular mechanisms that link wounding to reprogramming remain poorly understood. In this study we identify the Arabidopsis HEAT SHOCK FACTOR A1 (HSFA1) class of transcription factors, being key regulators of the heat stress response, as central players in wound-induced callus formation and shoot regeneration. Loss of HSFA1 function in the *hsfa1abd* triple mutants severely impairs cellular reprogramming, reducing callus formation from wounded hypocotyls, as well as shoot regeneration from explants. Conversely, overexpression of the HSFA1d gain-of-function variant markedly enhances regeneration. Time-series RNA-seq and ChIP-seq analyses revealed that HSFA1s directly activate the key reprogramming regulators *WOUND-INDUCED DEDIFFERENTIATION 1* (*WIND1*), *PLETHORA 3* (*PLT3*) and *ZINC FINGER OF ARABIDOPSIS THALIANA 6* (*ZAT6*). Furthermore, we demonstrate that HSFA1d activity is attenuated by SIZ1-mediated SUMOylation, linking post-translational modification to the regulation of wound responses. Our findings establish HSFA1s as an early transcriptional hub that integrates wound signals with the activation of a broad gene network that drives cellular reprogramming, thereby providing a mechanistic framework for understanding how stress-responsive transcription factors control regeneration.

## Introduction

Plants heal tissues and regenerate new organs upon amputation by reprogramming the developmental fate of somatic cells. Early wounding signals promote cellular reprogramming as do the phytohormones auxin and cytokinin that can be either endogenously produced or externally supplied. How these signals are perceived and integrated, however, remains a key focus of research in the field (Ikeuchi et al., 2020). One clear example where a plant’s regenerative capacity can be boosted by exogenous hormones is *de novo* shoot regeneration under an *in vitro* tissue culture condition. During this procedure an excised piece of tissue or explant is incubated on media containing auxin and cytokinin (Feldmann and David Marks, 1986; Valvekens et al., 1988). Preincubation on auxin-rich callus-inducing media (CIM) promotes cellular reprogramming and formation of a cell mass called callus that is characterized by the expression of root meristem regulators such as *PLETHORA 3* (*PLT3*), *PLT5*, *PLT7* and *WUSCHEL-RELATED HOMEOBOX 5* (*WOX5*) (Sugimoto et al., 2010; Kareem et al., 2015). This callus contains pluripotent cells which, upon incubation on cytokinin-rich shoot inducing media (SIM), give rise to shoot meristems.

Since intact plants are less able to regenerate shoots in tissue culture (Iwase et al., 2015), it is apparent that beyond phytohormones contained in the media, wounding signals are needed to trigger cellular reprogramming. Major regulators of stress-induced cellular reprogramming are the APETALA2/ETHYLENE RESPONSE FACTOR (AP2/ERF) transcription factors (TFs) WOUND INDUCED DEDIFFERENTIATION 1 (WIND1) and its homologs WIND2, WIND3 and WIND4 (Iwase et al., 2011; Iwase et al., 2015; Iwase et al., 2017; Iwase et al., 2018). Plants expressing a chimeric repressor of *WIND1*, i.e., *WIND1-SRDX,* display severely reduced shoot regeneration (Iwase et al., 2017), whereas ectopic overexpression of *WIND1* can overcome the need for wounding to induce shoot regeneration in tissue culture (Iwase et al., 2015). WIND1 promotes shoot regeneration by inducing the expression of the *ENHANCER OF SHOOT REGENERATION* (*ESR1)* gene (Iwase et al., 2017) as well as other genes required for callus formation and/or pluripotency acquisition (Iwase et al., 2021). Several other AP2/ERF TFs also play key roles in regeneration and for instance, ERF114 and ERF115 function together with their binding partners, SCARECROW-LIKE 5 (SCL5), SCL21 and PAT1 to promote cell proliferation, cell reprogramming and acquisition of stem cell fate upon wounding (Heyman et al., 2013; Heyman et al., 2016; Bisht et al., 2023). In addition, a number of pluripotency regulators, including *PLT3*, *PLT5*, *PLT7* and *WOX5*, are transcriptionally induced upon wounding (Ikeuchi et al., 2017; Rymen et al., 2019; Radhakrishnan et al., 2020). These factors are necessary for acquiring shoot fate, as loss-of-function mutants of *plt3 plt5 plt7* and *wox5 wox7 wox14* are deficient in shoot regeneration (Kareem et al., 2015; Kim et al., 2018). Despite the identification of these key transcriptional regulators of reprogramming, the upstream factors that link wound stress perception to the initiation of transcription remain largely unknown. A recent study in tomato reported that the REGENERATION FACTOR 1 (REF) peptide serves as a local wound signal that activates the *WIND1* expression, although the detailed underlying molecular mechanism is yet to be established (Yang et a., 2024).

HEAT SHOCK FACTORs (HSFs) are a family of transcription factors that play a central role in the response to heat stress (Andrasi et al., 2021). Among all HSFs, HSFA1, which in Arabidopsis is encoded by four homologs *HSFA1a*, *HSFA1b*, *HSFA1d* and *HSFA1e*, functions at the top tier of the heat-induced transcriptional network to induce the expression of many downstream genes, including other HSF homologs as well as *HEAT SHOCK PROTEINs* (*HSPs)* upon exposure to heat stress (Yoshida et al., 2011; Liu and Charng, 2013; Ohama et al., 2017; Li et al., 2019; Peng et al., 2025). Previous studies revealed that many *HSFs* and *HSPs* are also upregulated in response to a variety of non-heat related stresses including high light intensity, wounding, high salinity, drought and pathogen infection (Miller and Mittler, 2006; Swindell et al., 2007; Liu et al., 2011). The expression profiles of *HSF*s and *HSP*s are qualitatively similar between these stresses, although the timing of their expression may differ (Swindell et al., 2007). These observations suggest that HSFA1 and their downstream network play wider roles in the plant’s responses to biotic and abiotic stresses, but their exact functions outside of the heat stress response are not well understood.

All *HSFA1* genes are expressed prior to the perception of heat stress, and HSFA1 activity is controlled by the post-translational regulatory processes including those which involve HSFA1-HSPs interactions (Ohama et al., 2016). Under normal conditions HSP70/90 bind the temperature-dependent repression (TDR) domain of HSFA1, rendering HSFA1 inactive in the cytosol. Upon exposure to heat stress, HSFA1 proteins are released from HSP70/90 and become nuclear localised, where they activate the transcription of downstream genes (Ohama et al., 2016). A recent study further revealed that prion-like domains help concentrating HSFA1 into speckles after heat stress to enhance their transcriptional activity under such conditions (Peng et al., 2025). In addition, several lines of evidence suggest that HSFA1 might be modified by SMALL UBIQUITIN-LIKE MODIFIER (SUMO) through a process called SUMOylation. SUMOylation is a multi-step process in which the final conjugation step is catalysed by SAP & MIZ1 (SIZ1), one of only two E3 ligases in Arabidopsis (Miura et al., 2005; Ishida et al., 2009; Roy and Sadanandom, 2021). SUMOylation alters the activity of a number of important transcriptional regulators (Castro et al., 2012; Augustine and Vierstra, 2018) including several HSFs in Arabidopsis such as HSFA9, HSFA4a (Carranco et al., 2017), HSFA2 (Cohen-Peer et al., 2010) as well as SlHSFA1 in tomato (Zhang et al., 2018) and TaHSFA1 in wheat (Wang et al., 2023). Previous proteomics studies identified Arabidopsis HSFA1d as a potential SUMO target (Miller et al., 2010), but any functional relevance of this modification remains unknown.

In this study we demonstrate that wounding stress activates a HSFA1-regulated transcriptional cascade, promoting cellular reprogramming in Arabidopsis. Accordingly, our data show that HSFA1 proteins accumulate in the nucleus following wounding to induce the expression of key cellular reprogramming genes to promote callus formation and subsequent shoot regeneration. We further demonstrate that HSFA1d can be modified by SIZ1-mediated SUMOylation, leading to the repression of its transcriptional activity. This study thus reveals that the HSFA1 TFs function as early transducers of wound signalling in plant regeneration, with their activity being fine-tuned by post-translational mechanisms.

## Results

### Wounding promotes rapid accumulation of HSFA1 in nuclei to induce its target gene expression

To investigate how early transcriptional responses cause cellular reprogramming following wounding, we first identified genes that are rapidly induced upon wounding under a condition where Arabidopsis roots or hypocotyls naturally develop callus from cut sites without additional plant hormones. By reanalysing the RNA sequencing (RNA-seq) data from our previous studies (Ikeuchi et al., 2017; Rymen et al., 2019), we found that a total of 5615 genes were upregulated either within 1 or 3 hours of wounding in hypocotyls (n = 3882) or within 1, 3 or 12 hours in roots (n = 3665) (Supplementary Data Set 1). Gene ontology (GO) analysis surprisingly revealed that in addition to the terms such as “response to wounding”, the term “response to heat” was statistically enriched among the wound-induced genes in both hypocotyls (*P* = 1.32e-26) and roots (*P* = 7.07e-07) (Supplementary Data Set 1). This suggests that wounding induces the transcription of many genes implicated in the heat response. To test this hypothesis, we compared genes that are upregulated by wounding (n = 5615) with those upregulated in response to acute heat stress (n = 6024) (Cortijo et al., 2017; Ikeuchi et al., 2017; Li et al., 2019; Rymen et al., 2019). We found that 34% of the genes induced by wounding were also induced by heat stress (Fig. 1A; Supplementary Data Set 1, *P* = 7.4e-13). Previous studies have shown that HSFA1 is required for the induction of 355 genes after heat shock (Liu and Charng, 2013; Li et al., 2019). Interestingly, around two thirds, i.e., 208 genes out of 355 genes, among these heat-induced HSFA1 target genes were also upregulated by wounding (Fig. 1B; Supplementary Data Set 1, *P* = 1.5e-32). In particular, among the 32 heat-induced HSFA1 target genes that encode TFs, 26 genes are upregulated by wounding in roots and/or hypocotyls (Fig. 1C). As reported for heat response (Liu and Charng, 2013; Friedrich et al., 2021), the transcript level of most of these TF genes are transiently upregulated within 1 to 3 hours after wounding and later return to essentially basal levels of expression (Fig. 1C). Importantly, several members of the HSF family genes, including *HSFA3*, *HSFA7A*, *HSFB2A*, *HSF4/HSFB1* and *HSFA2*, are included among these heat/wounding-induced TF genes. This suggests that wounding activates the canonical HSFA1-mediated transcriptional network. Several other TF-encoding genes, such as *DEHYDRATION-RESPONSIVE ELEMENT-BINDING PROTEIN 2A* (*DREB2A*), *ZINC FINGER OF ARABIDOPSIS THALIANA* (*ZAT6*) and *WIND1* known to function in various abiotic responses such as drought and oxidative stress (Sakuma et al., 2006; Iwase et al., 2011; Chen et al., 2016; Tang and Luo, 2018), are also included among these genes, further highlighting the common transcriptional activation between these abiotic stresses.

**Figure 1.**
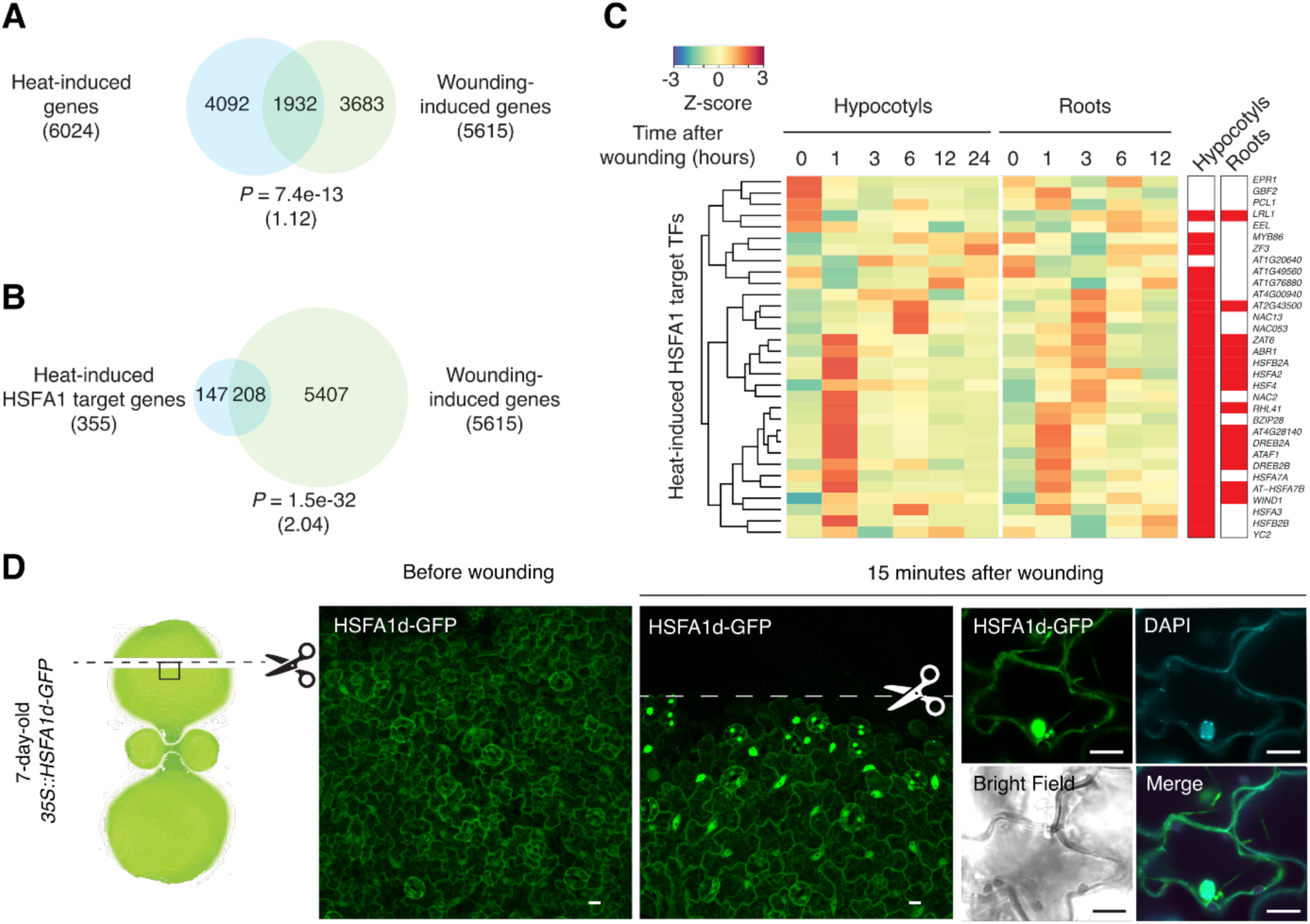
Wounding promotes translocation of HSFA1 to nuclei. (A) Venn diagram of heat-induced genes and wound-induced genes. Heat-induced genes include those upregulated within 30 minutes (Li *et al*, 2019) or 4 hours after incubation at 37°C (Cortijo *et al*, 2017b). Wound-induced genes include those upregulated at 1, 3, 6 or 12 hours in roots (Rymen *et al*, 2019) or at 1 or 3 hours in hypocotyls after wounding (Ikeuchi *et al*, 2017). The significance of the overlap between the pair of genes was evaluated by a hypergeometric test. The calculated representation factor is shown in a bracket. (B) Venn diagram of genes induced by heat stress in the HSFA1 dependent manner (Liu & Charng, 2013; Li *et al*, 2019) and those induced by wounding in Fig. 1A. The significance of the overlap between the pair of genes was evaluated by a hypergeometric test. The calculated representation factor is shown in a bracket. (C) Heatmap showing the expression of heat-induced HSFA1 target TFs (Liu & Charng, 2013; Li *et al*, 2019) after wounding in hypocotyls (Ikeuchi *et al*, 2017) or roots (Rymen *et al*, 2019). Genes upregulated at least at one time point are marked in red in the rightmost sidebars. (D) A diagram showing the set-up for confocal observation of wound-induced protein localization (left) and confocal images showing the subcellular localisation of HSFA1d-GFP proteins before and 15 minutes after wounding in the cotyledon epidermis of 1 week-old *35S::HSFA1d-GFP* seedlings (middle). Close-up images confirming the localisation of HSFA1d-GFP proteins in DAPI-staining nuclei (right). Scale bars, 10 µm.

Given that heat shock or warm temperatures induces the nuclear localisation of HSFA1d (Yoshida et al., 2011; Ohama et al., 2016; Li et al., 2024; Peng et al., 2025), we tested whether wounding affects the localisation of HSFA1 proteins. As reported previously (Yoshida et al., 2011; Ohama et al., 2016), the majority of HSFA1d-GFP proteins are found within the cytosol of intact tissue expressing *HSFA1d-GFP* under the control of constitutive *Cauliflower Mosaic Virus 35S* promoter (*35S::HSFA1d-GFP*) (Fig. 1D). In contrast, HSFA1d-GFP proteins accumulate in the nucleus within 15 minutes following wounding (Fig. 1D). A similar nuclear localisation of HSFA1d-GFP could be also observed following wounding in the *pHSFA1d::HSFA1d-GFP hsfa1abd* line in which HSFA1d-GFP complements the thermotolerance phenotype in *hsfa1abd* (Ohama et al., 2016) (Supplementary Fig. S1). These results demonstrate that HSFA1d rapidly relocates from the cytoplasm to the nuclei in response to wounding, thereby activating its downstream transcriptional network.

### HSFA1 TFs are required for cellular reprogramming during regeneration

One of the typical wound responses displayed by many plants is the formation of callus at the wound sites (Iwase et al., 2011; Kareem et al., 2025). To investigate the function of HSFA1 in this wound response, we first tested whether they are involved in callus formation of wounded hypocotyls (Fig. 2A). Among the four HSFA1 members in Arabidopsis the combined mutation in three of these, i.e. HSAF1a, HSFA1b, and HSFA1d, severely impairs thermotolerance (Yoshida et al., 2011). As the *hsfa1abd* mutants have a mixed genetic background of Columbia (Col-0) and Wassilewskija (Ws), of which the wild type (WT) plants display different rates of callus formation following wounding (Supplementary Fig. S2A), we compared the phenotypes of *hsfa1abd* versus *pHSFA1d::HSFA1d-GFP hsfa1abd* plants (Ohama et al., 2016). We found that the *hsfa1abd* mutants make callus statistically less frequently compared to *pHSFA1d::HSFA1d-GFP hsfa1abd* plants (Supplementary Fig. S2A), illustrating that HSFA1 is required for callus formation in response to wounding. To validate the involvement of HSFAs in this process, we used CRISPR-Cas9 to generate two triple mutants (*hsfa1abd-1* and *hsfa1abd-2*) in the WT Col-0 background (Supplementary Fig. S2, C and D). Additionally, we crossed *hsfa1abd-2* with the *hsfa1e* (Col-0) to generate the *hsfa1abde* quadruple knockout mutants. As shown in Fig. 2, A and B, both the *hsfa1abd-1* and *hsfa1abd-2* triple mutants form callus less frequently than WT and the quadruple mutants exhibit an even greater impairment in callus formation. Furthermore, while WT and *hsfa1d-1* display higher callus formation rates at 25°C compared to 22°C, the higher-order mutants regenerate less callus at the warmer temperature, where HSFA1 is expected to be more active (Cortijo et al., 2017; Dickinson et al., 2018) (Fig. 2, A and B). These data demonstrate that the HSFA1 TFs are required for callus formation in response to wounding.

**Figure 2.**
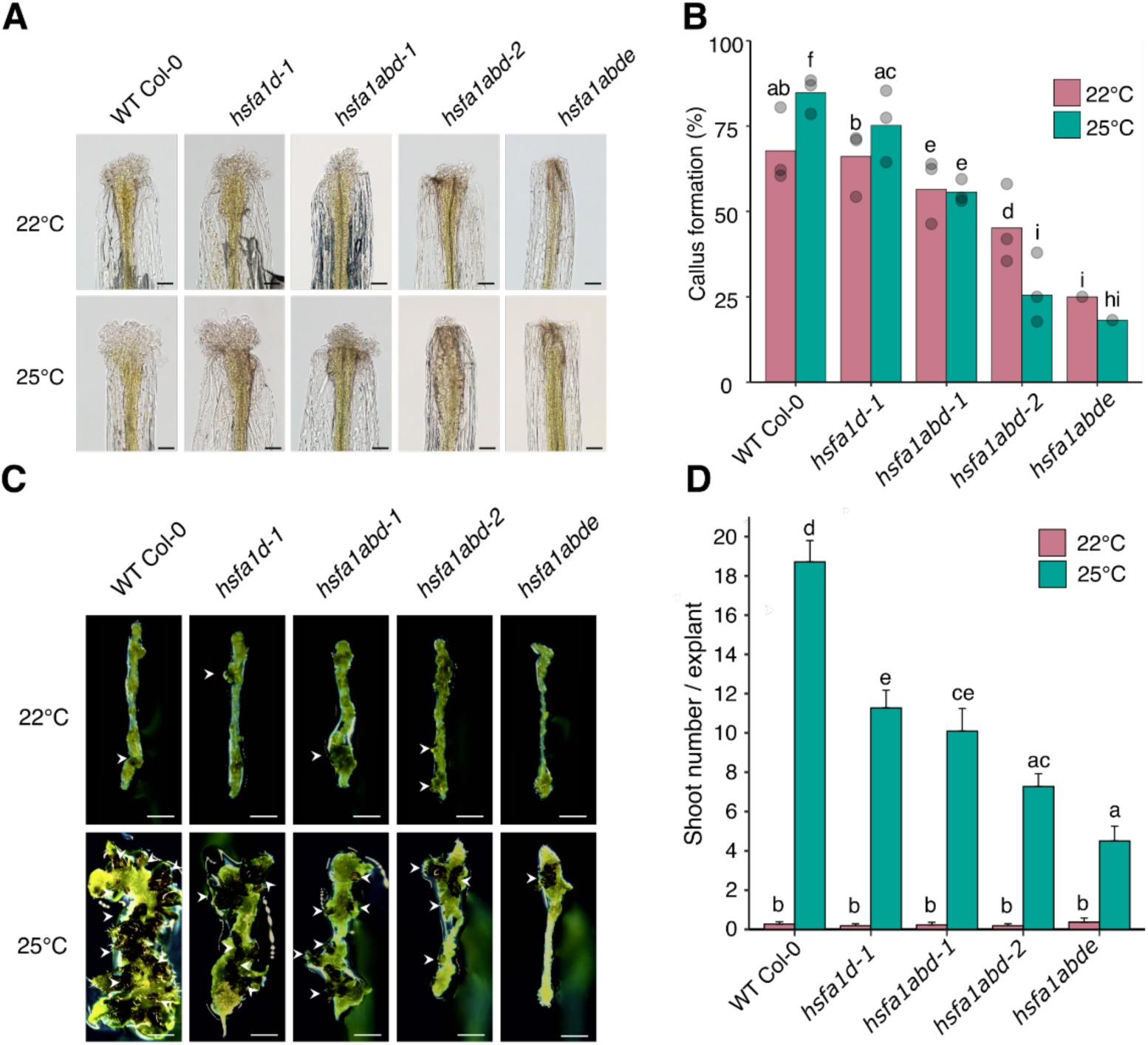
HSFA1 TFs are required for cellular reprogramming in regeneration. (A) Images of callus formation from the wound site of WT Col-0, *hsfa1d-1*, *hsfa1abd-1, hsfa1abd-2* and *hsfa1abde* hypocotyls. Scale bars = 100 µm. (B) Quantification of the proportion hypocotyls from (A) that form callus within one week after cutting (right). Sample sizes are WT Col-0 (22°C *n* = 121, 25°C *n =* 131), *hsfa1d-1* (22°C *n* = 115, 25°C *n =* 117), *hsfa1abd-1* (22°C *n* = 108, 25°C *n =* 106)*, hsfa1d-2* (22°C *n* = 93, 25°C *n =* 98) and *hsfa1abde* (22°C *n* = 24, 25°C *n =* 22). Individual points are the callus-formation rates from three independent experiments. Different letters indicate significant differences based on pairwise two-proportions z-test (*P* < 0.05). (C) Images of shoot regeneration from hypocotyl explants of WT Col-0, *hsfa1d-1*, *hsfa1abd-1, hsfa1d-2* and *hsfa1abde*. All explants were incubated on CIM for 4 days and then on SIM for 14 days at 22°C or 25°C. Arrows indicate regenerated shoots. Scale bars, 2 mm. (D) Barplot showing the quantification of shoot regeneration from hypocotyls in (C). Values represent mean number of shoots produced per explant, and error bars represent ± SE. Sample sizes are WT Col-0 (22°C *n* = 22, 25°C *n =* 21), *hsfa1d-1* (22°C *n* = 22, 25°C *n =* 22), *hsfa1abd-1* (22°C *n* = 22, 25°C *n =* 22)*, hsfa1d-2* (22°C *n* = 22, 25°C *n =* 22) and *hsfa1abde* (22°C *n* = 22, 25°C *n =* 22). Different letters indicate significant differences based on one-way ANOVA and posthoc Tukey test (*P* < 0.05).

Given that wounding-induced cues are vital for changing cell fate in organ regeneration (Iwase et al., 2015; Ikeuchi et al., 2019; Ikeuchi et al., 2020), we next tested the role of HSFA1 in the context of *in vitro* shoot regeneration. We compared the HSFA1 mutants to WT using the two-step CIM and SIM conditions (Valvekens et al., 1988). As previously reported (Lambolez et al., 2022), hypocotyl explants regenerated significantly more shoots at 25°C than at 22°C (Fig. 2, C and D). Moreover, we found that all HSFA1 mutants, including *hsfa1d-1,* regenerated significantly fewer shoots compared to WT or *pHSFA1d::HSFA1d-GFP hsfa1abd-2* explants at 25°C (Fig. 2, C and D; Supplementary Fig. S2E). The quadruple mutant *hsfa1abde* explants regenerated the fewest shoots under these conditions, with around a third of the number produced by WT (Fig. 2C). Importantly, all of these *hsfa1* mutant explants are capable of developing callus in the presence of exogenous plant hormones on CIM but their callus appears to be less efficient at gaining the shoot fate on SIM (Fig. 2C), suggesting that HSFA1 is required for the pluripotency acquisition and/or subsequent cell fate conversion. Similar phenotypes were observed for the *hsfa1abd* explants in the mixed Col-0/Ws background that showed significantly reduced shoot regeneration compared to the *pHSFA1d::HSFA1d-GFP hsfa1abd* explants (Supplementary Fig. S2B). These data thus demonstrate that the HSFA1 TFs are required for cellular reprogramming during wounding-induced callus formation and *in vitro* shoot regeneration.

### Hyperactivation of HSFA1 enhances cellular reprogramming

Overexpression of *HSFA1* TFs and consequential hyperactivation of their targets confer tolerance to several abiotic stresses (Ogawa et al., 2007; Bechtold et al., 2013; Ohama et al., 2016; Tian et al., 2020). To investigate whether *HSFA1* overexpression enhances cellular reprogramming, we examined the regeneration phenotype of *35S::HSFA1d-GFP* plants. As HSFA1d activity can be repressed by HSP70/HSP90 when they bind the TDR domain within the HSFA1d protein, we also examined the *35S::HSFA1dΔ1-GFP* line with deleted TDR domain, previously shown to exhibit a constitutively active heat stress response (Ohama et al., 2016). Although the *35S::HSFA1dΔ1-GFP* plants show much lower mRNA and protein expression levels compared to *35S::HSFA1d-GFP* (Supplementary Fig. S3A), overexpression of *HSFA1dΔ1-GFP*, but not of *HSFA1d-GFP*, significantly promotes callus formation in wounded hypocotyls (Fig. 3A). Consistently, while plants overexpressing the full-length *HSFA1d-GFP* regenerate a similar number of shoots as the WT, *35S::HSFA1dΔ1-GFP* plants exhibit a drastically increased number of regenerated shoots (Fig. 3B). This supports the idea that increased HSFA1 activity boosts cellular reprogramming in regeneration.

**Figure 3.**
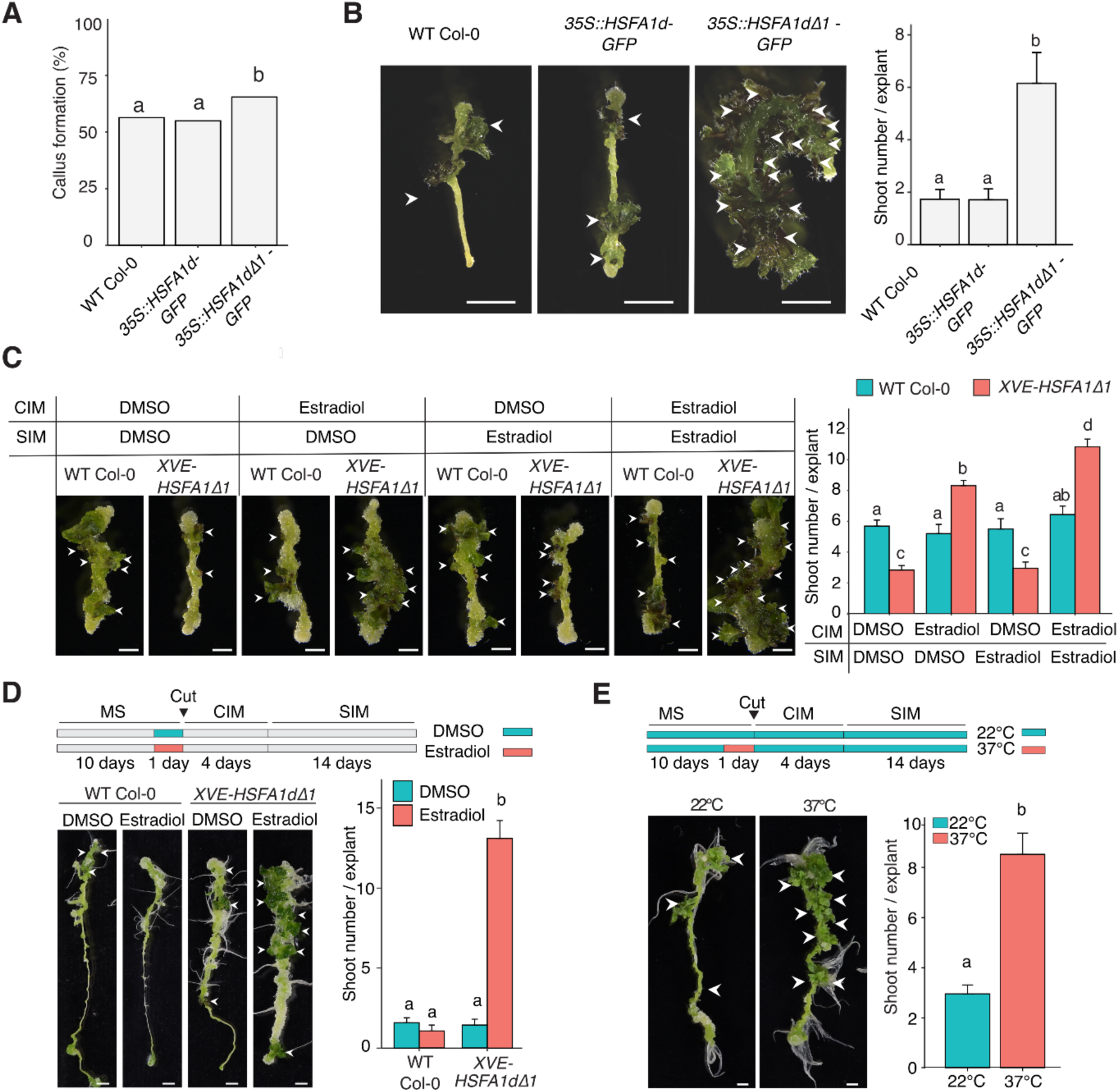
Hyperactivation of HSFA1 promotes shoot regeneration. (A) Quantification of the proportion of hypocotyl explants that form callus within one week after cutting. Sample sizes are WT Col-0 (*n* = 103), *35S::HSFA1d-GFP* (*n* = 122), and *3S::HSFA1dΔ1-GFP* (*n* = 81). Different letters indicate significant differences based on pairwise two-proportions z-test (*P* < 0.05). (B) Images (left) and quantification (right) of shoot regeneration from hypocotyl explants of WT Col-0, *35S::HSFA1d-GFP* and *35S::HSFA1dΔ1-GFP*. Sample sizes are WT Col-0 (*n* = 54), *35S::HSFA1d-GFP* (*n* = 73) and *35S::HSFA1dΔ1-GFP* (*n* = 73). Values represent mean number of shoots produced per explant, and error bars represent ± SE. Different letters indicate significant differences based on one-way ANOVA and posthoc Tukey test (*P* < 0.05). Scale bars, 2 mm. (C) Representative images (left) and quantification (right) of shoot regeneration from WT Col-0 and *XVE-HSFA1dΔ1* hypocotyl explants. Explants were incubated on CIM for 4 days followed by SIM for 14 days containing either DMSO or 1 µM 17-ß-estradiol as indicated. Sample sizes are WT Col-0 treated with DMSO (*n* = 26), WT Col-0 treated with 17-ß-estradiol (*n* = 16-20), *XVE-HSFA1dΔ1* treated with DMSO (*n* = 48) and *XVE-HSFA1dΔ1* treated with 17-ß-estradiol (*n* = 31-42). Values represent mean number of shoots produced per explant, and error bars represent ± SE. Different letters indicate significant differences based on one-way ANOVA and posthoc Tukey test (*P* < 0.05). Scale bars, 2 mm. (D) A diagram showing the experimental setup for the transient *HSFA1dΔ1* induction (top). Representative images (left) and quantification (right) of shoot regeneration from WT Col-0 and *XVE-HSFA1dΔ1* root explants incubated in the presence of DMSO or 1 µM 17-ß-estradiol. Sample sizes are WT Col-0 treated with DMSO (*n* = 12), WT Col-0 treated with 17-ß-estradiol (*n* = 17), *XVE-HSFA1dΔ1* treated with DMSO (*n* = 21) and *XVE-HSFA1dΔ1* treated with 17-ß-estradiol (*n* = 19). Values represent mean number of shoots produced per explant, and error bars represent ± SE. Different letters indicate significant differences based on one-way ANOVA and posthoc Tukey test (*P* < 0.05). Scale bars, 2 mm. (E) A diagram showing the experimental setup for the heat stress treatment (top). Representative images (left) and quantification (right) of shoot regeneration from WT Col-0 root explants. Sample sizes are 22°C (*n* = 57) and 37°C (*n* = 55). Values represent mean number of shoots produced per explant, and error bars represent ± SE. Different letters indicate significant differences based on one-way ANOVA and posthoc Tukey test (*P* < 0.05). Scale bars, 2 mm.

We next aimed to determine the stage at which *HSFA1dΔ1* overexpression promotes shoot regeneration. To this end, we generated *XVE-HSFA1dΔ1* plants that express *HSFA1dΔ1* under the control of the estradiol-inducible XVE system (Zuo et al., 2000). By RT-qPCR we verified that application of 1 µM 17-β-estradiol to these transgenic plants increased the *HSFA1d* transcript level by up to 280 times compared to the mock (DMSO) treatment within 20 hours of treatment (Supplementary Fig. S3B). Using these inducible lines, we observed that *HSFA1dΔ1* overexpression in hypocotyl explants during either culturing on CIM, or during culturing on both CIM and SIM, enhances shoot regeneration (Fig. 3C). Transient overexpression of *HSFA1dΔ1* for 24 hours before wounding, however, does not enhance shoot regeneration in hypocotyl explants (Supplementary Fig. S3C). These data show that maintaining high levels of HSFA1 activity following wounding on CIM helps shoot regeneration in hypocotyls.

To assess whether the HSFA1’s regeneration-promoting activity extends across tissues, we then examined whether *HSFA1dΔ1* overexpression can stimulate shoot regeneration from root explants. As with hypocotyls, we saw that roots constitutively overexpressing *HSFA1dΔ1* produce significantly more shoots than either WT or *35S::HSFA1d* (Supplementary Fig. S3D). Notably, however, in contrast to hypocotyls, inducing *HSFA1dΔ1* just 24 hours before root explant excision enhanced shoot regeneration when compared to both WT and DMSO-treated *XVE-HSFA1dΔ1* explants (Fig. 3D). These data demonstrate that HSFA1 class of TFs promotes regeneration in a tissue context-dependent manner, with its effects in roots being particularly pronounced when induced near the time of wounding.

Based on these observations, we next asked whether heat stress itself could promote shoot regeneration. We incubated WT seedlings at 37°C for 24 hours immediately before explant cutting and incubation on CIM and SIM. While hypocotyl explants show no enhancement in shoot regeneration (Supplementary Fig. S3E), heat-treated root explants produce significantly more shoots on SIM than those kept at 22°C (Fig. 3E), clearly showing that transient hyperactivation of heat response prior to wounding is sufficient to enhance shoot regeneration.

### HSFA1 TFs are required for the transcriptional activation of reprogramming associated genes after wounding

Having established the central role of the HSFA1 TFs in cellular reprogramming, we next sought to investigate the underlying molecular mechanisms. To this end, we first carried out a time-series RNA-seq analysis by using WT and *hsfa1abd-2* to capture the transcriptomic changes within 24 hours after wounding (Fig. 4A). At T0, 828 genes were already downregulated in the mutant, suggesting that HSFA1 regulates the expression of many genes not associated with stress response. This was increased to 1404 and 1607 genes at 1 hour and 24 hours following wounding, respectively (Supplementary Fig. S4A and Data Set 2), indicating that HSFA1 also activates a large number of wound-responsive genes. Consistently, GO analysis of these downregulated genes revealed an enrichment for categories such as “response to heat”, “response to oxidative stress” and “response to wounding” are enriched (Supplementary Fig. S4B). Because HSFA1 functions at the top tier of a multi-layered transcriptional network (Yoshida et al., 2011; Liu and Charng, 2013; Ohama et al., 2017; Li et al., 2019; Peng et al., 2025), we focused on the 241 TFs that were downregulated in wounded *hsfa1abd-2* hypocotyls at one or more time points (Fig. 4B). Many of these genes are rapidly induced within 30 minutes to 1 hour following wounding in WT but this early activation is strongly reduced in the mutant. Another distinct TF cluster is also induced in an HSFA1-dependent manner later at 24 hours (Fig. 4B).

**Figure 4.**
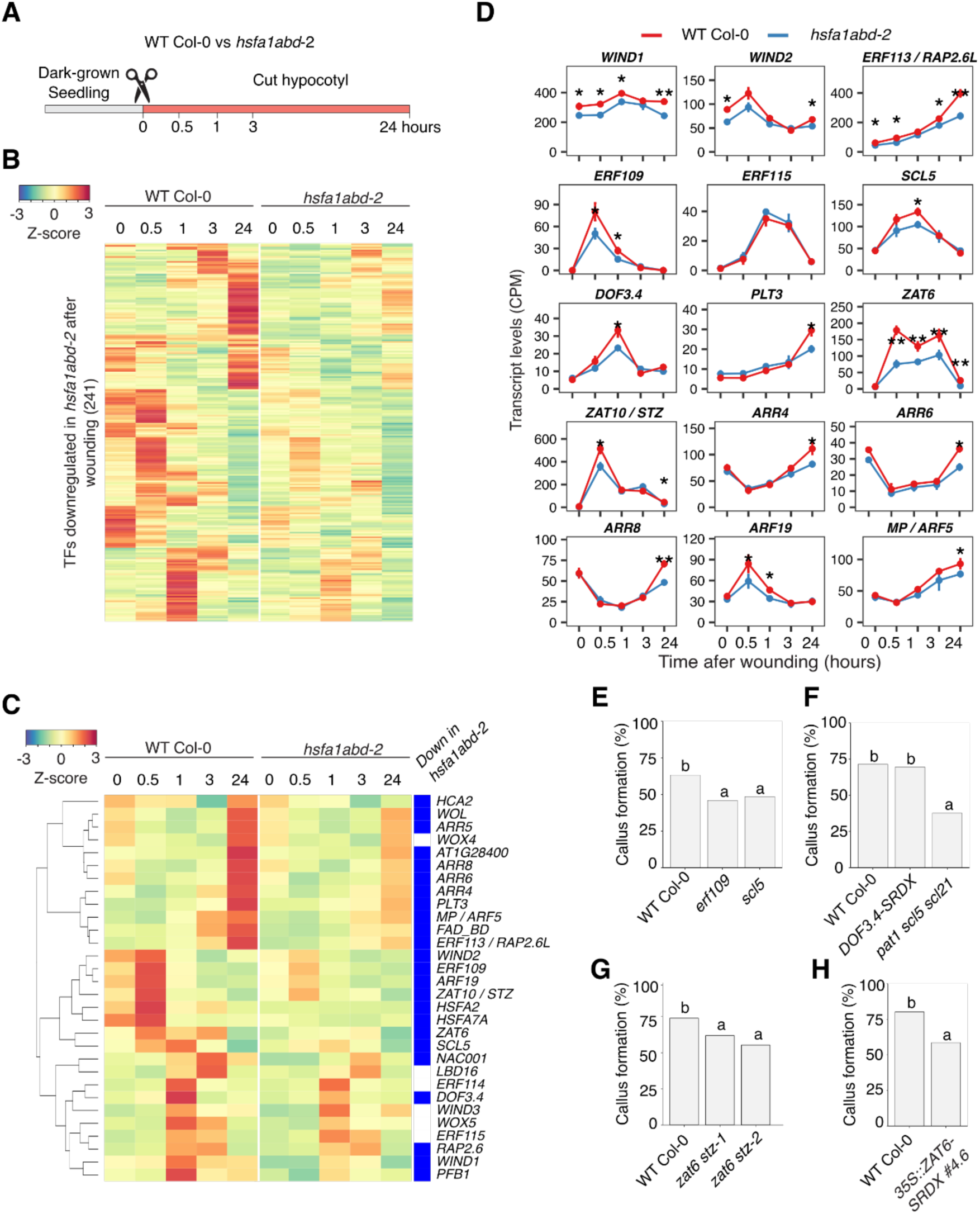
HSFA1 TFs are required for the transcriptional upregulation of reprogramming associated genes after wounding. (A) A diagram showing the RNA-seq experiment setup. WT Col-0 and *hsfa1abd-2* seedlings grown in the dark at 22°C were wounded and hypocotyls were harvested at 0, 0.5, 1, 3, and 24 hours after wounding for RNA extraction and sequencing. (B) Heatmap showing the expression of 241 transcription factors that were downregulated at any timepoint in *hsfa1abd-2* compared to WT Col-0. (C) Heatmap showing the normalised expression of selected genes related to regeneration or known HSFA1 target genes. The green sidebar indicates genes that are bound by HSFA1-GFP based on ChIP-seq data. The blue sidebar indicates genes that are significantly downregulated in *hsfa1abd-2* at any time point after wounding based on RNA-seq. (D) Line plot showing transcript levels (CPM) of selected genes in (A). Error bars represent SE and asterisks indicate statistical differences based in Edge-R analysis (* *P* < 0.05, ** *P* < 0.01). (E-H) Quantification of the proportion of hypocotyl explants that form callus within one week after cutting. Sample sizes are: (E) WT Col-0 (*n* = 141), *erf109* (*n* = 109), and *scl5* (*n* = 202); (F) WT Col-0 (*n* = 72), *DOF3.4-SRDX* (*n* = 132), and *pat1 scl5 scl21* (*n* = 56); (G) WT Col-0 (*n* = 142), *zat6 stz-1* (*n* = 105), and *zat6 stz-2* (*n* = 103); (H) WT Col-0 (*n* = 136) and *35S::ZAT6-SRDX #4.6* (*n* = 82). Different letters indicate significant differences based on pairwise two-proportions z-test (*P* < 0.05).

Among the HSFA1-regulated TFs, we identified a subset of genes previously implicated in stress-induced cellular reprogramming. The expression of *WIND1* and *WIND2*, for instance, is significantly reduced in *hsfa1abd-2* both before and after wounding, suggesting that HSFA1 regulates their expression during normal development and in response to injury (Fig. 4, C and D). Among other AP2/ERF family members involved in wound-induced reprogramming (Heyman et al., 2013; Heyman et al., 2016; Ikeuchi et al., 2017; Zhang et al., 2019), *ERF113* and *ERF109* expression is acutely induced by wounding in an HSFA1-dependent manner (Fig. 4, C and D). While *ERF115* expression is not affected by HSFA1, we did notice that its binding partner *SCL5* and a downstream target of both HSFA1 and SCL5, *DNA-BINDING ONE FINGER 3.4* (*DOF3.4*), are downregulated in *hsfa1abd-2* (Fig. 4, C and D). Previous studies have shown that the *PLT* genes are also wound-induced regulators of callus formation (Ikeuchi et al., 2017; Iwase et al., 2021) and our data demonstrate that the expression is also regulated in an HSFA1-dependent manner (Fig. 4, C and D). Consistent with earlier work showing that *WIND1-SRDX* plants and *plt* mutants display defects in callus formation in wounded hypocotyls (Iwase et al., 2011; Ikeuchi et al., 2017), we found that *erf109, scl5,* and higher order *pat1 scl5 scl21* mutants (Bisht et al., 2023) also form callus less frequently compared to WT (Fig. 4, E and F). However, wounded hypocotyls from dominant negative *35S::DOF3.4-SRDX* plants (Bisht et al., 2023) show no obvious defects in callus formation (Fig. 4F), suggesting that the ERF115-SCL5 pathway regulates callus formation through mechanisms independent of DOF3.4.

Since the putative targets of HSFA1 include many TFs that have not been previously characterized in the context of the wound response, we further examined whether some of these contribute to regeneration. Among these, the C2H2 zinc-finger transcription factors ZAT6 and ZAT10, also known as SALT TOLERANCE ZINC FINGER (STZ), are particularly interesting since they are both highly wound-inducible and their expression depends on HSFA1 (Fig. 1C, 4C and 4D). We thus generated two independent double knock-out mutants of ZAT6 and STZ using CRISPR-Cas9 (Supplementary Fig. 5A) and assessed callus formation from wounded hypocotyls. As shown in Fig. 4G, *both zat6 stz-1 and zat6 stz-2* have impaired callus formation (Fig. 4G). Consistently, expressing the dominant negative ZAT6-SRDX proteins causes impaired callus formation in *35S::ZAT6-SRDX* (Fig. 4H), strongly suggesting that ZAT6 functions as a regulator of wound-induced callus formation.

Notably, several HSFA1-regulated TFs detected in this study are known to respond to auxin or cytokinin (Fig. 4, C and D). Consistent with previous observation that wounding is accompanied with elevated cytokinin signalling (Iwase et al., 2011; Ikeuchi et al., 2017), the cytokinin-responsive A-type response regulators *ARABIDOPSIS RESPONSE REGULATOR4* (*ARR4*)*, ARR5, ARR6* and *ARR8* are upregulated within 24 hours after wounding in WT and this is dependent on HSFA1 (Fig. 4, C and D). Furthermore, the auxin-responsive *AUXIN RESPONSE FACTOR 19* (*ARF19*) is rapidly and transiently upregulated at 30 minutes after wounding but its expression is reduced in *hsfa1abd-2* (Fig. 4, C and D). In contrast, *MONOPTEROS/ARF5* expression increases more gradually, becoming evident at 3 hours, yet it too requires HSFA1 for full induction (Fig. 4, C and D). Together, these results show that HSFA1 activates a broad array of wound-or hormone-responsive transcriptional regulators that collectively contribute to wound-induced cellular reprogramming.

### HSFA1 TFs directly induce *WIND1*, *PLT3* and *ZAT6* to promote cellular reprogramming

In order to reveal which TF genes are directly regulated by HSFA1, we subsequently performed chromatin immunoprecipitation-sequencing (ChIP-seq) analysis using the *pHSFA1d::HSFA1d-GFP hsfa1abd* after heat stress or wounding. As reported by Peng et al. (2025), subjecting seedlings to heat stress dramatically increases binding of HSFA1d-GFP to target DNA sequences (Fig. 5A; Supplementary Data Set 3). Similarly, wounding induces HSFA1d binding within 10 minutes and a large portion of these go back to the basal levels after 3 hours (Fig. 5A; Supplementary Data Set 3), implying the relatively transient nature of HSFA1-mediated transcriptional regulation. As expected, the loci bound by HSFA1d-GFP upon heat and wounding treatment largely overlap, with 1175 of the 2365 genes bound after heat stress also bound after wounding in at least one of our time points (Supplementary Data Set 3), supporting our view that these stresses regulate at least a partly shared set of genes. By comparing the 320 HSFA1d-bound TF genes (Supplementary Data Set 3) with the 241 TF genes downregulated in *hsfa1abd-2*, we found that 33% TF genes (81 out of 241) in common (Fig. 5B, Supplementary Data Set 4), suggesting that HSFA1d directly upregulates these target genes. Importantly, these analyses confirmed the previously reported direct targets of HSAF1, such as *HSFA2*, *HSFA3*, *HSFA7A*, *HSFB1* and *HSFB2A*, and in addition identified several key regulators of reprogramming including *WIND1*, *WIND2*, *PLT3, ZAT6, STZ* and *SCL5* as directly targeted by HSFA1 (Supplementary Data Set 4).

**Figure 5.**
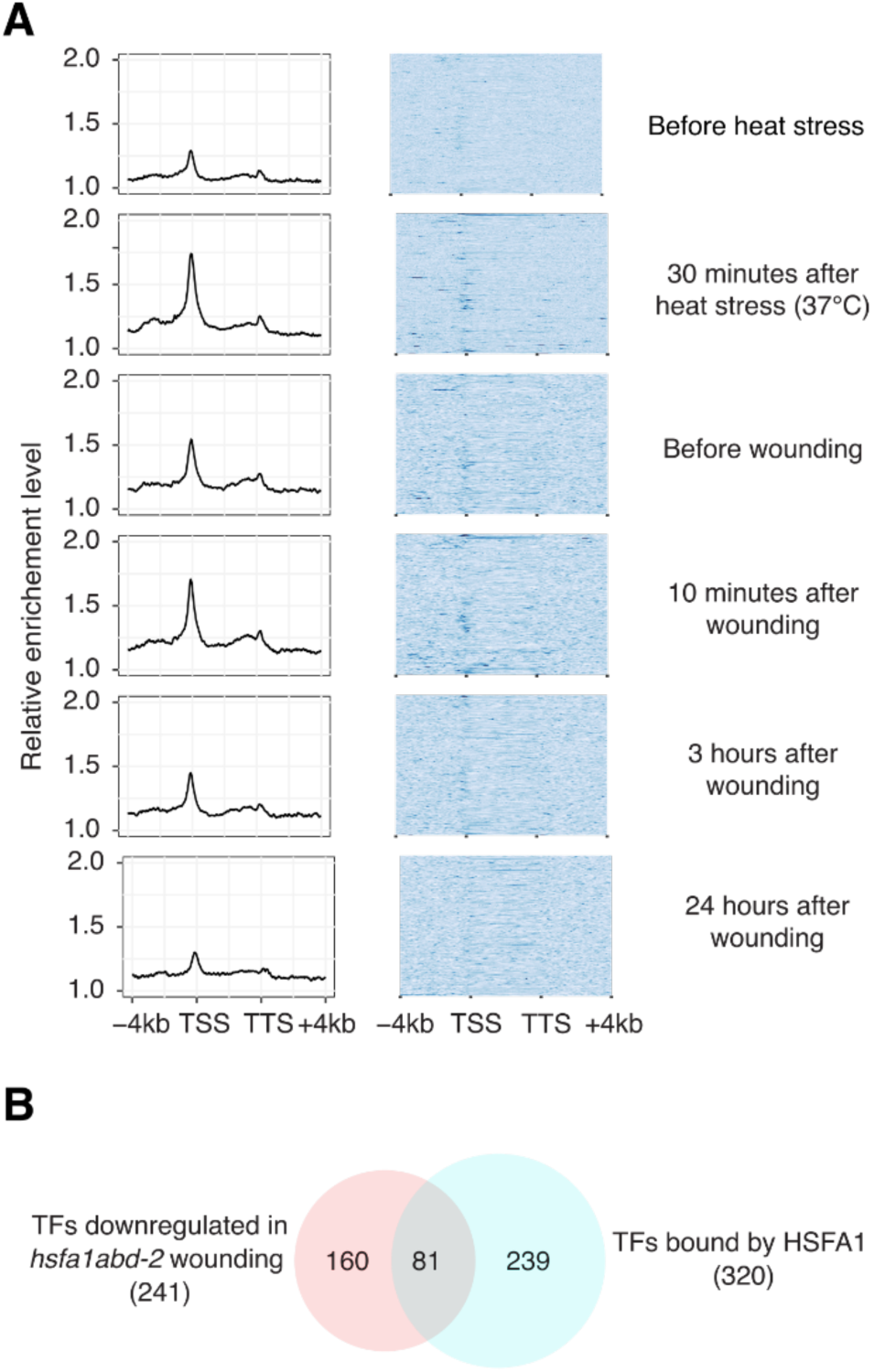
HSFA1 TFs directly activate a large set of genes after wounding. (A) Line plot (left) and heatmap (right) showing normalized HSFA1d-GFP enrichment at genes bound by HSFA1d-GFP at 22°C, after 30 minutes at 37°C or after 10 minutes, 3 hours and 24 hours after wounding based on ChIP-seq analysis. Values indicate binding signals normalized to the input. (B) Venn diagram showing the overlap of transcription factors downregulated in *hsfa1abd-2* (Fig. 4B) and bound by HSFA1d-GFP (Fig. 5A).

Among these HSFA1d direct target genes, we first focused our attention to WIND1 since wounding increases the binding of HSFA1d-GFP to the *WIND1* locus (Fig. 6A; Supplementary Data Set 3). By performing an electrophoretic mobility shift assay (EMSA) with purified MBP-HSFA1d proteins, we verified the binding of HSFA1d to the promoter region of *WIND1* that contains a cis-regulatory motif known as heat shock element (HSE)-like element (GAGAGTTC) (Fig. 6B). In addition, we observed that this binding is abolished by either adding a non-labelled competitor probe or mutating the HSE-like sequence (Fig. 6B), indicating that the detected binding is sequence specific. We have previously reported that the expression of the dominant-negative *WIND1-SRDX,* severely inhibits shoot regeneration (Iwase et al., 2017). Notably, introduction of *WIND1-SRDX* into *35S::HSFA1dΔ1* plants strongly suppresses the enhanced shoot regeneration phenotype (Fig. 6C; Supplementary Fig. S5B), supporting that WIND1 functions downstream of HSFA1.

**Figure 6.**
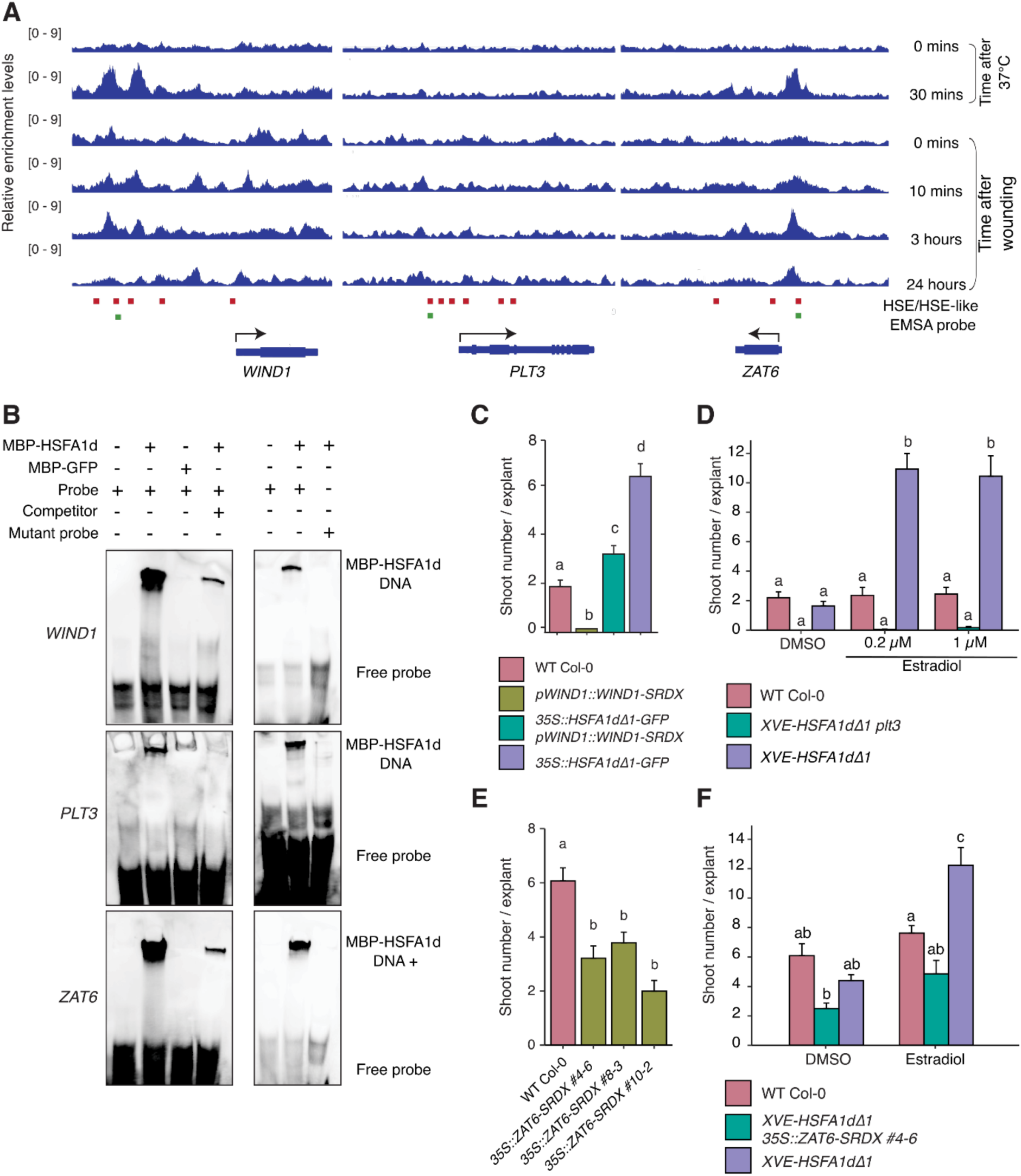
HSFA1 TFs directly activate the expression of *WIND1*, *PLT3* and *ZAT6* to promote cellular reprogramming. (A) Integrative Genomics Viewer traces of HSFA1d-GFP enrichment at *WIND1*, *PLT3* and *ZAT6* locus at 22°C, after 30 minutes at 37°C or after 10 minutes, 3 hours and 24 hours after wounding based on ChIP-seq data (Fig. 5A). Arrows indicate the directionality of transcription. Red boxes mark the position of HSE or HSE-like elements and green boxes highlight the position of electrophoretic mobility shift assay (EMSA) probes used in Fig. 6B. (B) EMSA showing the *in vitro* binding of MBP-HSFA1d proteins to the cis elements of *WIND1*, *PLT3* and *ZAT6* genes. Recombinant MBP-HSFA1d proteins or MBP-GFP proteins were incubated with 40-bp biotin-labelled DNA probes. Probe sequences were determined based on ChIP-seq data and non-labelled probes were used as competitors (left). Putative binding motifs were mutated to generate mutant probes (right) (C) Quantification of shoot regeneration from hypocotyl explants of WT Col-0, *pWIND1::WIND1-SRDX*, *35S::HSFA1dΔ1-GFP pWIND1::WIND1-SRDX,* and *35S::HSFA1dΔ1-GFP*. All explants were incubated on CIM for 4 days and then on SIM for 14 days. Values represent mean number of shoots produced per explant, and error bars represent ± SE. Sample sizes are WT Col-0 (*n* = 36), *pWIND1::WIND1-SRDX* (*n* = 26), *35S::HSFA1dΔ1-GFP pWIND1::WIND1-SRDX* (*n* = 42) and *35S::HSFA1dΔ1-GFP* (*n* = 39). Different letters indicate significant differences based on one-way ANOVA and posthoc Tukey test (*P* < 0.05). (D) Quantification of shoot regeneration from hypocotyl explants of WT Col-0, *XVE-HSFA1dΔ1 plt3* and *XVE-HSFA1dΔ1*. Explants were incubated on CIM and SIM containing DMSO, 0.2 µM or 1 µM 17-ß-estradiol. Values represent mean number of shoots produced per explant, and error bars represent ± SE. Sample sizes are 30 explants for all conditions. Different letters indicate significant differences based on one-way ANOVA and posthoc Tukey test (*P* < 0.05). (E) Quantification of shoot regeneration from hypocotyl explants of WT Col-0 and *35S::ZAT6-SRDX.* Sample sizes are WT Col-0 (*n* = 46), *35S::ZAT6-SRDX* (middle; *n* = 35-42). Different letters indicate significant differences based on one-way ANOVA and posthoc Tukey test (*P* < 0.05). (F) Quantification of shoot regeneration from hypocotyl explants of WT Col-0*, XVE::HSFA1dΔ1 35S::ZAT6-SRDX and XVE-HSFA1dΔ1* explants were incubated on CIM and SIM containing either DMSO or 1 µM 17-ß-estradiol. Values represent mean number of shoots produced per explant, and error bars represent ± SE. Sample sizes are WT Col-0 (*n* = 12-16), *XVE-HSFA1dΔ1 35S::ZAT6-SRDX* (*n* = 21-26) and *XVE-HSFA1dΔ1* (*n* = 15-22). Different letters indicate significant differences based on one-way ANOVA and posthoc Tukey test (*P* < 0.05).

ChIP-seq analysis, in addition, showed that HSFA1d binds the promoter of *PLT3* near multiple HSEs, and this binding was confirmed *in vitro* by EMSA (Fig. 6, A and B; Supplementary Data Set 3). To assess if PLT3 acts downstream of HSFA1, we introduced the *plt3* mutation into *XVE-HSFA1dΔ1*. As expected, the *plt3* mutation completely abolished the ability for HSFA1dΔ1 to promote shoot regeneration (Fig. 6D; Supplementary Fig. S5C). Lastly, we tested whether ZAT6 functions downstream of HSFA1 in regeneration since our ChIPseq and EMSA data show that HSFA1d binds the HSE element of its promoter upon wounding (Fig. 6, A and B, Supplementary Data Set 3). As shown in Fig. 6E, F and Supplementary Fig. S5B, C, both *zat6 stz* double mutants and *35S::ZAT6-SRDX* explants regenerate fewer shoots than WT, and introduction of *35S::ZAT6-SRDX* into *XVE-HSFA1dΔ1* compromised shoot regeneration, supporting that ZAT6 also promotes shoot regeneration downstream of HSFA1. These observations together establish that HSFA1 promotes shoot regeneration by directly activating the expression of several key reprogramming regulators.

### SIZ1-mediated SUMOylation prevents hyperactivation of the HSFA1d activity

We previously demonstrated that the SUMO E3 ligase SIZ1 negatively regulates shoot regeneration by repressing wound-induced gene expression (Coleman et al., 2020). An earlier proteomics study, in addition, suggested that HSFA1d can be SUMOylated after heat shock (Miller et al., 2010), thus it is possible that SIZ1 SUMOylates HSFA1d to attenuate its transcriptional activity. To validate this hypothesis, we performed an immunoprecipitation experiment using whole proteins extracted from *35S::HSFA1d-GFP* plants. As heat strongly induces protein SUMOylation (Kurepa et al., 2003; Cohen-Peer et al., 2010), we subjected these seedlings to heat stress by incubating at 37°C for 1 hour. After pulling down the HSFA1d-GFP proteins with antibodies against GFP, we probed these proteins with the antibodies against Arabidopsis SUMO1 (Ishida et al., 2009) to detect SUMO conjugation. As shown in Fig. 7A, we observed that the full-length HSFA1d protein is highly SUMOylated after heat stress treatment. Interestingly, we also found that the SUMOylation of HSFA1dΔ1-GFP is reduced to around 16% of the levels observed in full-length HSFA1d protein (Fig. 7A). These data confirm that HSFA1d is SUMOylated after heat shock and further show that the 36 amino acids truncated in the HSFA1dΔ1 protein are required for full heat shock-induced SUMOylation of the protein.

**Figure 7.**
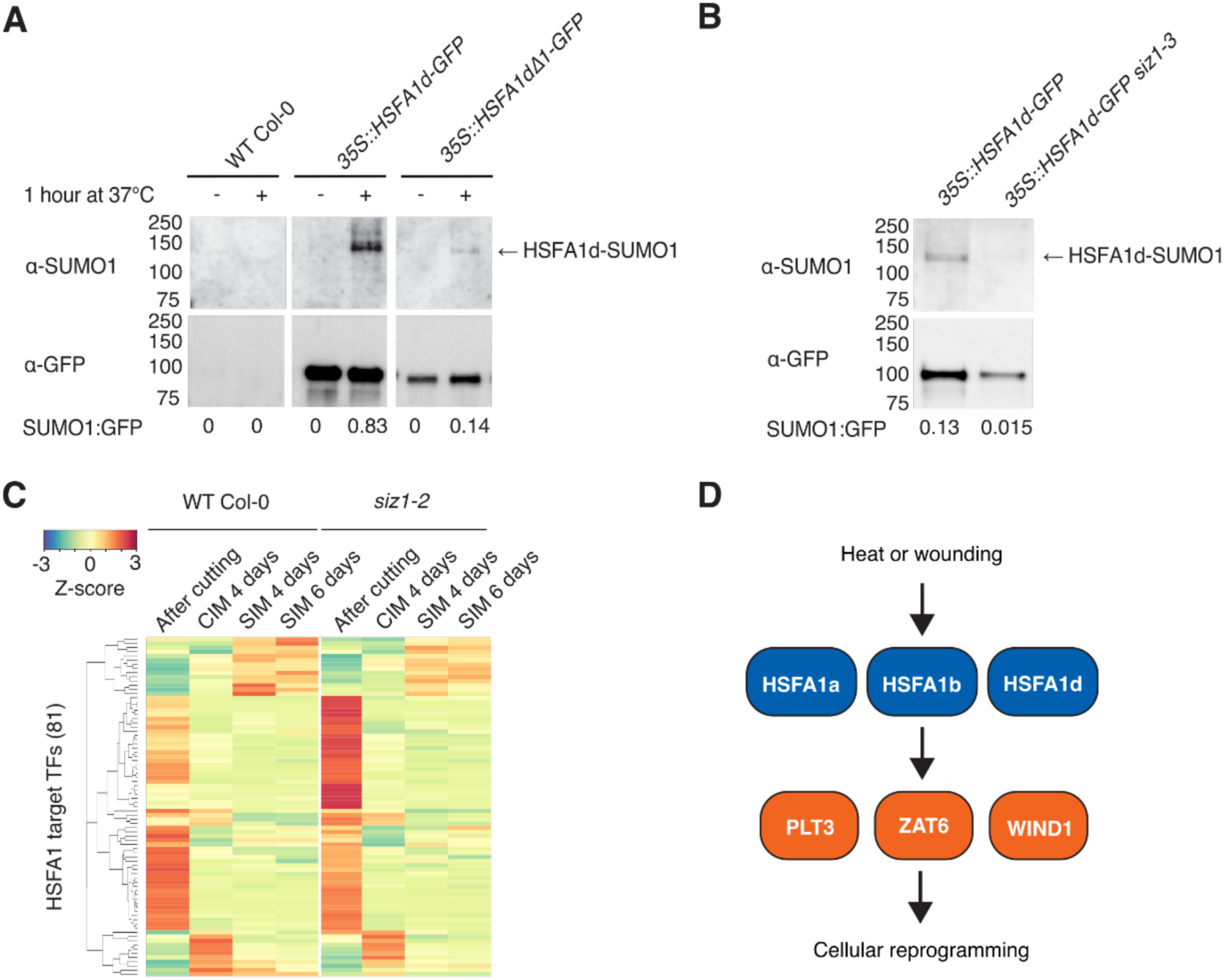
The HSFA1 activity is regulated by SIZ1-mediated SUMOylation. (A) Immunodetection of heat-induced SUMOylation on HSFA1d-GFP proteins. For heat treatment, WT Col-0, *35S::HSFA1d-GFP* and *35S::HSFA1dΔ1-GFP* seedlings were incubated at 37°C for 1 hour. HSFA1d-GFP proteins were immunoprecipitated with anti-GFP antibodies. HSFA1-GFP proteins and their SUMOylation were detected by anti-GFP antibodies and anti-SUMO1 antibodies, respectively. These bands were quantified with ImageJ and their relative values are shown. (B) Immunodetection of heat-induced SUMOylation on HSFA1d-GFP proteins in *siz1-3*. For heat treatment, WT Col-0, *35S::HSFA1d-GFP* and *35S::HSFA1-GFP siz1-3* seedlings were incubated at 37°C for 1 hour. HSFA1d-GFP proteins were immunoprecipitated with anti-GFP antibodies. HSFA1d-GFP proteins and their SUMOylation were detected by anti-GFP antibodies and anti-SUMO1 antibodies, respectively. These bands were quantified with ImageJ and their relative values are shown. (C) Heatmap showing the expression of heat-induced HSFA1 target TFs in WT Col-0 and *siz1-2* after cutting, CIM 4 days, SIM 4 days or SIM 6 days. Data are from Coleman *et al*, (2020). (D) A schematic model showing the HSFA1-mediated transcriptional cascade acting after wounding stress. Wounding rapidly promotes nuclear translocation of HSFA1 and it then directly activates *WIND1, PLT3* and *ZAT6* expression to promote cellular reprograming.

To examine if the SUMOylation of HSFA1d is dependent on SIZ1, we introduced the *35S::HSFA1d-GFP* construct into the *siz1-3* mutant background and performed the same immunoprecipitation experiment. As expected, the *siz1-3* mutation causes strong reduction of the HSFA1d-SUMO1 signal (Fig. 7B), demonstrating that SIZ1 plays a key role in SUMOylation of HSFA1d. To test if the HSFA1 transcriptional activity is altered in the *siz1* mutants, we analysed the expression of heat-induced HSFA1 target TF genes using RNA-seq data we previously published (Coleman et al., 2020). As shown in Fig. 7C, many of these HSFA1 targets, such as other HSFs and DREBs, are transiently induced after cutting in WT and their expression is strongly increased in *siz1-2*. Importantly, the expression of *WIND1* and *ZAT6,* which displays transient induction upon wounding in WT, is significantly upregulated in *siz1-2* (Coleman et al., 2020) (Fig. 7C). These data thus support that HSFA1d is a key target of SUMOylation in the context of regeneration and SUMOylation prevents hyperactivation of its target genes that include core reprogramming regulators we identified in this study.

## Discussion

This study revealed that the HSFA1 TFs are activated upon wounding and they function as key regulators of cellular reprogramming in wound-induced callus formation and shoot regeneration. Our data show that HSFA1 rapidly accumulates within nuclei after wounding and promotes reprogramming by directly inducing the expression of *WIND1*, *PLT3*, and *ZAT6* (Fig. 7D). Furthermore, we identify SIZ1-mediated SUMOylation as a probable negative regulatory mechanism that attenuates HSFA1 activity, likely preventing hyperactivation of the HSFA1 pathway under stress conditions. This regulatory balance ensures robust but controlled activation of wound-induced transcriptional programmes, integrating injury signals with developmental responses.

### Roles of HSFA1 TFs in the wounding response

We show that wounding triggers the nuclear localisation of HSFA1d within minutes, coinciding with the transcriptional activation of its downstream genes (Fig. 1C, 1D and 4C). Previous studies have hinted at such a role by noting that many HSPs and HSFs are upregulated by abiotic stresses including wounding (Miller and Mittler, 2006; Swindell et al., 2007). Our work provides direct evidence that HSFA1d is activated by wounding and is essential for the wound-induced expression of its target genes.

What signals promote the activation of HSFA1d in response to wounding remains unknown but insights from heat stress studies may provide some clues. For instance, HSFA1 proteins can directly sense heat stress via prion-like domains that facilitate transcriptionally-active protein speckle formation at target loci (Peng et al., 2025). It would be interesting to test if this also happens after wounding and whether such enhancer elements exist near the HSFA1 target genes identified in this study. Additionally, heat-induced calcium influx is thought to activate HSFA1 by the calmodulin-binding protein kinase 3 (CBK3)-dependent phosphorylation (Liu et al., 2008; Saidi et al., 2009; Finka et al., 2012). Since wounding creates a rapid intracellular calcium influx (Mousavi et al., 2013), it would be worth investigating if increasing cellular calcium levels activates HSFA1 upon wounding or if sequestering calcium (e.g. EGTA treatment) attenuates its activity. Other potential activators of HSFA1 are reactive oxygen species (ROS), which also accumulate upon both heat and wounding stress (Miller et al., 2009; Beneloujaephajri et al., 2013; Prasad et al., 2019; Devireddy et al., 2021). It is reported that HSFA1a trimerisation and DNA binding are dependent on the redox state *in vitro* (Liu et al., 2013), and that the nuclear translocation of HSFs may depend on ROS (Giesguth et al., 2015). Further work is required to clarify whether HSFA1 is regulated by wounding-induced ROS signalling. Besides these stress-induced signals, brassinosteroid (BR) is recently shown to activate HSFA1 since under non-stressed conditions, nuclear translocation of HSFA1d is suppressed by BR-INSENSITIVE 2 (BIN2)-mediated phosphorylation (Luo et al., 2022). Heat and BR dephosphorylate HSFA1d and permit its nuclear translocation to initiate the target gene expression. Although an involvement of BR in wound-induced processes is not established, this is an interesting possibility that should be investigated in future studies. Finally, HSFA1 is negatively regulated by HSP70/90 binding, while the interaction is abolished in response to heat stress (Ohama et al., 2016). It is likely that wounding also relieves the HSP70/90 repression since removing the HSP interacting TDR domain enhances wound-induced callus formation and shoot regeneration (Fig. 3B; Supplementary Fig.S2 B).

In addition, we show that SIZ1-dependent SUMOylation attenuates the wound-induced expression of HSFA1 targets (Fig. 7C). Although we have not established an experimental system to detect SUMOylation after wounding, we predict that wounding induces SUMOylation of HSFA1d like heat treatment to safeguard cells against hyperactivation. This is supported by the observation of the strongly reduced SUMOylation in the *35S::HSFA1dΔ1* plant combined with the hyper-regeneration of this same mutant (Fig. 3B and 7A). SUMOylation is known to repress the transactivation activity of the mammalian HSFA1 homolog HSF1 (Hietakangas et al., 2006), implying that this mechanism is conserved between animals and plants. Our data show that SUMOylation is strongly reduced in *35S::HSFA1dΔ1-GFP* plants (Fig. 7A), thus it will be interesting to test if SUMOylation participates in the regulation of HSFA1 binding to HSP70/90. Although previous reports are conflicting as to whether HSFA1 target gene expression is changed in Arabidopsis *siz1* mutants (Yoo et al., 2006; Catala et al., 2007), our data clearly show that SIZ1 negatively regulates the expression of HSFA1 downstream targets (Fig. 7C). HSFA1 SUMOylation is likely part of the complex post-translational regulatory processes that fine-tune the HSFA1 activity, and future research should investigate exactly how these are coordinated to ensure its fast and transient activation in response to wounding.

### Roles of HSFA1 in cellular reprogramming

It is well established that HSFA1 helps plants survive severe stresses by producing protective chaperone proteins and ROS scavenging enzymes. In animals, HSFs are also known to function in developmental processes (Akerfelt et al., 2010) and emerging evidence started to reveal roles of HSFA1 and other HSF homologs in plant development (Pernas et al., 2010; ten Hove et al., 2010; Liu and Charng, 2013; Wunderlich et al., 2014; Albihlal et al., 2018). Our findings expand this view by showing that HSFA1 integrates wounding responses and developmental programmes, enabling plants to cope with injury by promoting wound-healing callus formation and regeneration of severed organs. HSFA1 appears to act as one of the earliest wounding-activated regulators of cellular reprogramming described to date, representing the top tier of the wound-induced transcriptional network (Fig. 7D).

Our data show that once activated, HSFA1 binds the promoter of key reprogramming regulators, including *WIND1*, *PLT3* and *ZAT6*, to induce their expression (Fig. 4, C and D, Fig. 6, A and B). This finding is consistent with a previous report showing that *WIND1* and *ZAT6* induction by heat stress is compromised in the *hsfa1abde* quadruple mutants (Li et al., 2019). Our results, in addition, suggest that HSFA1 activates a broader network by inducing multiple other AP2/ERF TFs and their partners both directly and indirectly (Fig. 4, C and D; Supplementary Data Set 4), thereby amplifying the AP2/ERF-mediated transcriptional programmes. Notably, we identified the C2H2 zinc-finger TF ZAT6 as a previously unrecognized regulator of callus formation and shoot regeneration (Fig. 4, G and H, Fig. 6E). Future work should explore how the ZAT6-mediated pathway interfaces with other previously identified regulators such as WIND1 and PLTs to coordinate downstream gene expression during reprogramming. We also note that the induction of these HSFA1 downstream genes is not completely abolished in the *hsfa1abd* mutant (Fig. 4, C and D). This could be because HSFA1e, still functional in the *hsfa1abd* mutant, can activate their expression and/or the HSFA1 pathway functions in parallel with other wound-activated pathways. In the context of root regeneration from detached leaves, for instance, jasmonate-induced pathway regulates the expression of reprogramming genes such as *ERF109* (Zhang et al., 2019). It is thus possible that wounding signalling is transduced through the multiple signalling pathways to orchestrate the activation of the reprogramming gene regulatory network.

It is intriguing that wounding induces several auxin– and cytokinin-responsive genes in an HSFA1-dependent manner (Fig. 4, C and D), suggesting that HSFA1 may modulate these hormonal signalling pathways in reprogramming. Notably, induction of four A-type ARRs clearly depends on HSFA1, and our ChIP-seq data suggest that some of these are direct HSFA1 targets (Fig. 4, C and D; Supplementary Data Set 4). This raises the possibility that certain A-type ARRs transcriptionally respond not only to elevated cytokinin but also directly to wounding stress itself. As A-type ARRs act as negative regulators of cytokinin signalling, such wound-induced activation may serve as an additional safeguard to fine-tune the cytokinin response during reprogramming. Although evidence from a previous study indicates that auxin signalling is not important for callus formation in wounded hypocotyls (Ikeuchi et al., 2017), we observed HSFA1-dependent induction of *MP/ARF5* and *ARF19*. Determining whether these ARFs contribute to reprogramming, and whether they respond to auxin and/or other wounding-derived cues will be an important direction for future studies.

In conclusion, our findings uncover the HSFA1 family as a central regulator that connects early wound-induced signals to the transcriptional networks driving cellular reprogramming. This work broadens the roles of the HSFA1 family beyond previously known abiotic stress responses and establishes it as a pivotal component of the wound-induced regeneration programme in plants.

## Materials and Methods

### Plant materials and growth conditions

The Columbia (Col-0) and Wassilewskija (Ws) ecotypes of *Arabidopsis thaliana* (Arabidopsis) were used. The mixed background *hsfa1abd* mutant was previously described (Yoshida et al., 2011). To generate *hsfa1abd-1* and *hsfa1abd-2* triple mutants, CRISPR-Cas9-mediated gene editing was done in *hsfa1d-1* (SALK_022404) (Liu et al., 2011) plants using the pKAMA-ITACHI vectors (Tsutsui and Higashiyama, 2017) and gRNAs targeting HSFA1a and HSFA1b (Supplementary Data Set 5). The *hsfa1abde* quadruple mutant was generated by crossing *hsfa1abd-2* with *hsfa1e* (SALK_094943) (Liu et al., 2011).). To generate *zat6* and *stz* mutants, gRNAs targeting both genes were cloned into *pFASTRK-pRPS5A-BP-AtCas9-BP*-G7T-A-BsaI-ccdB-BsaI-G* by golden-gate cloning as described (Develtere et al., 2024) and homozygous single and double mutants were selected by PCR with primers in Supplementary Data Set 5. The transgenic line *pHSFA1d::HSFA1d-GFP hsfa1abd-2* was generated by transforming *hsfa1abd-2* with *pHSFA1d::HSFA1d-GFP* constructs. The transgenic lines *pHSFA1d::HSFA1d-GFP hsfa1abd*, *35S::HSFA1d-GFP* and *35S:HSFA1dΔ1-GFP* (Lohmann et al., 2004; Yoshida et al., 2011; Ohama et al., 2016), and *WIND1-SRDX* (Iwase et al., 2011) were previously described. Homozygous plants for T-DNA mutants were selected by PCR using primers listed in Supplementary Data Set 5. Seeds were stratified for 3 days at 4°C and then sown onto Murashige and Skoog (MS) media containing 1% Sucrose and 0.6% (w/v) Gelzan^™^ (Sigma-Aldrich). Seedlings were incubated under constant fluorescent white light (approximately 50 µmol m^-2^ s^-1^) at 22°C.

### Cloning of *HSFA1dΔ1* and generation of *XVE-HSFA1dΔ1* plants

Genomic DNA was extracted from plants carrying the *35S::HSFA1dΔ1* transgene (Ohama et al., 2016) and the coding region of *HSFA1dΔ1* was PCR amplified with primers listed in Supplementary Data Set 5, cloned into the entry vector pENTR/D-TOPO (Invitrogen) and then subcloned into the gateway-compatible binary vector pER8GW (Papdi et al., 2008) using the Gateway LR Clonas II Enzyme (Invitrogen). The binary vector was transformed into WT Col-0 plants using the floral-dip method (Clough and Bent, 1998). Seedlings containing the transgene were selected on MS media containing 50 mg ml^-1^ spectinomycin and 50 mg ml^-1^ hygromycin. Homozygous T3 plants were selected by PCR genotyping.

### *in vitro* shoot regeneration assay

Seedlings were grown in the dark for 7 days to induce etiolation. Hypocotyl explants (around 10 mm long) were excised using a razor blade (Feather) and incubated on CIM (Gamborg B5 medium with 0.25% Gelzan, 0.5 μg/ml 2,4-dichlorophenoxyacetic acid [2,4-D] and 0.05 μg/ml kinetin) for 4 days under constant light. Hypocotyl explants were then transferred to SIM (Gamborg B5 medium with 0.25% Gelzan, 0.15 µg/ml IAA and 0.5 µg/ml 2-IP) and incubated for several days to induce shoot regeneration. To quantify shoot regeneration on SIM, the number of shoots visible per explant was counted. Shoots were defined as regions with viable leaves with trichomes and appearing to arise from a single meristem as visualized from the top with an OLYMPUS SZX7 microscope.

### Wound-induced callus formation assay

Wound-induced callus assay was performed by cutting the hypocotyl of 7-day-old dark-grown seedlings about 3 mm below the shoot apical meristem and incubating them on MS in constant light at 22°C for 4 days. Presence of callus was assessed by formation of more than 3 callus cells protruding from the cut site when visualized with an Olympus SZX7 microscope.

### Confocal microscopy

Plants expressing *35S::HSFA1d-GFP* were grown in constant light at 22°C for 7 days. Plants were mounted in water before imaging the adaxial side of the cotyledon with a SP8 confocal microscope equipped with a 63X water immersion objective lens (Leica Microsystems). GFP was excited by a 488 nm laser obtained from a highly flexible pulsed white-light laser and measured with an emission spectrum of 498–580 nm using a hybrid detector with time gating of autofluorescence as previously described (Kodama, 2016). DAPI was excited by a 405 nm laser and measured with an emission spectrum of 412–476 nm using a hybrid detector. To wound the explant, the cotyledon was cut with micro scissors, remounted in water and incubated under light for 15 minutes. A z-stack was taken through the sample, and projected images were generated using the ImageJ Bio-Formats plugin with maximum of slices option.

### RNA extraction and quantitative RT-qPCR

RT-qPCR was performed as previously described (Shibata et al., 2018). Total RNA was extracted from 7-day-old whole seedlings using a RNeasy Plant Mini Kit (Qiagen). Extracted RNA was reverse transcribed using a PrimeScript RT-PCR kit with DNase I (Perfect Real Time) (Takara) in accordance with the accompanying protocol. Transcript levels were determined by RT-qPCR using a THUNDERBIRD SYBR qPCR Mix kit (Toyobo) and an Mx399P QPCR system (Agilent). For each sample, the expression levels were quantified for 3 biological replicates and normalized to those of the *PP2A3* gene. All primer sequences are listed in Supplementary Data Set 5.

### RNA-seq analysis

WT and *hsfa1abd-2* seedlings were grown in the dark for 7 days before their hypocotyls were excised and wounded as described in Ikeuchi et al., (2017), around 50 explants were collected per replicate and three biological replicates were used for each time point as described in Coleman et al. (2020). RNA was isolated using the RNeasy plant mini kit (Qiagen) and then subjected to library preparation with the Kapa stranded mRNA sequencing kit (KK8420, Kapa Biosystems) and Illumina-compatible FastGene adapters (NGSAD24, Nippon Genetics). Paired-end sequencing was performed on a NovaSeqX platform, and read trimming was carried out using fastp (Chen et al., 2018) with the following settings; –q 15 –l 5 –w 16 –x. Processed reads were mapped to the TAIR10 genome with STAR (Dobin et al., 2013) with the following settings; ––quantMode GeneCounts ––outFilterMultimapNmax 1 ––alignIntronMax 10000 alignMatesGapMax 10000.

Genes with the P value < 0.05 using the edgeR package (Robinson et al., 2010) in R were considered as differentially expressed and expression level was normalized with the cpm function in the edgeR package. Genes with Volcano plots were made using the ggplot2 package (Villanueva and Chen, 2019) and clusterProfiler (Yu et al., 2012) was used for GO analysis. The GO enrichment data were simplified using the “simplify” function and redundant GO categories, referring to the same biological functions, were further removed manually. The heatmaps.2 function in the gplots package was used to generate heatmaps. Venn diagrams were generated using the Vennerable package (https://r-forge.r-project.org/projects/vennerable) in R was used.

### ChIP-seq analysis

For heat-stress treatment, three biological replicates of approximately 60 2-week-old *pHSFA1d::HSFA1d-GFP hsfa1abd* seedlings grown at 22°C were subject to heat stress by transferring the plates to a 37°C incubator for 30 minutes. For wounding treatment, approximately 60 2-week-old seedlings grown at 22°C were cut multiple times with a razor blade and incubated on MS for 10 minutes, 3 hours or 24 hours. Plants were then frozen in liquid Nitrogen and ground using a bead shocker (MB1200 Yasui Kikai) in 50 ml tubes and the nuclear fraction was isolated after cross-linking for 10 min using 1% formaldehyde (Sigma) under vacuum. Chromatin was sheared at 4– 6 °C for 15 min with a focused ultrasonicator (Covaris) with the following settings: duty cycle 5%, intensity 4, and cycles per burst 200. Sheared chromatin was immunoprecipitated using antibodies against GFP (ab290, Abcam) and input DNA was kept as a control. The isolated DNA was quantified with the Qubit dsDNA High Sensitivity Assay kit (Thermo Fisher Scientific), and 1–5 ng of DNA was used to make each ChIP-seq library. Libraries were prepared using the KAPA Hyper Prep Kit for Illumina (KK8502, KAPA Biosystems) and Illumina compatible adaptors (E7335, E7500, E7710, E7730, NEB). The 300–500 bp DNA fragments were enriched using Agencourt AMPure XP (Beckman Coulter). Libraries were pooled and 50-bp, single-read sequences were obtained with Illumina NextSeq500 sequencer at a depth of at least 10 million mapped reads. Reads were mapped to the TAIR10 Arabidopsis genome using Bowtie2 (Langmead and Salzberg, 2012) with a mapping rate of 70 to 83%. SAM files were converted to BAM format using SAMTOOLS (Li et al., 2009). Mapped BAM files were subject to peak calling using MACS2 (Zhang et al., 2008) with a p-value cutoff of 0.05, m-fold of 5 50 and using the input sample as a control. Peaks were annotated to genes or their promoter (3 kb upstream of the TSS) using a custom bash script. To normalise the peak data, BAM files were indexed in SAMTOOLS and then normalized using bamCompare in deepTools2 (Ramirez et al., 2016) with the settings; –bs 2 ––operation ratio ––scaleFactorsMethod SES. To make Heatmaps, a matrix was generated with computeMatrix in deepTools2 with normalised BAM files. A Heatmap was then generated with ggplot2 geom_raster (Villanueva and Chen, 2019), 2016) following clustering with hclust in R. Tracks were also visualized in integrated genome viewer (IGV) browser (Thorvaldsdottir et al., 2013). Heat shock elements (HSE) and HSE-like motifs were defined as GAANNTTC, TCCNNGAA, TCNAGANNNNTC, WCNAGAANNNTC and GAGAGTTC (Bonner et al., 2000).

### Recombinant protein purification and electrophoresis mobility shift assay

The HSFA1d cDNA was amplified by PCR with primers listed in Supplementary Data Set 5. The fragment was purified and cloned into pENTR™⁄D-TOPO™ entry vector (ThermoFisher Scientific), followed by LR recombination with the destination vector pMGWA. To express the recombinant protein, E. coli (BL21-AI™) harbouring the pMGWA-HSFA1d vector were incubated with 0.4mM IPTG and 0.2% (w/v) L-arabinose at 37°C. Cells were harvested and lysed by sonication and the soluble fraction was kept for protein purification. MBP-HSFA1d was purified using the MBP-Spin Protein Miniprep Kit (Zymo Research) according to manufacturer’s instructions. EMSA assay was carried out using LightShift® Chemiluminescent EMSA Kit (Thermo Scientific). Purified proteins were incubated for 20 minutes at room temperature with the biotin-labelled DNA probe (175 fmol) or unlabelled DNA probe (28 pmol) in DNA-binding Buffer (10mM Tris, 50mM KCl, 1mM DTT, 5mM MgCl2, 0.05% NP-40, 2.5% glycerol, 0.05 µg.µl^-1^ Poly (dI•dC); pH 7.5) with a reaction volume of 20 µl. Sample was run on a native polyacrylamide electrophoresis gel, transferred and UV-crosslinked to a nylon membrane. Biotin-labelled DNA was detected by chemiluminescence with Streptavidin-Horseradish Peroxidase Conjugate.

### Immunoprecipitation and western blot analysis

Approximately 60 2-week-old seedlings grown at 22°C were subject to heat stress (37°C 1 hour), frozen in liquid Nitrogen and ground using a bead shocker (MB 1200 Yasui Kikai) in 50 ml tubes. Subsequently 4-5 ml of extraction buffer (100mM Tris-HCl (pH 8.0), 0.1% (w/v) SDS, 0.5% (v/v) Sodium Deoxycholate, 1% (v/v) Glycerol, 50mM Sodium Metabisulfite, 20mM N-Ethylmaleimide (NEM), cOmplete™ Protease Inhibitor Cocktail (Sigma-Aldrich), Pefabloc SC (Sigma-Aldrich) was added and mixed. Samples were then cleared by centrifugation at 2500 rpm 15 minutes and protein concentration of supernatant was measured by Bradford assay (Bio-Rad). Sample was diluted (1/10) in water and 10 μl was mixed with Bradford reagent in a 96 well plate and incubated at room temperature for 15 minutes. Absorbance was read using a plate reader (Infinite 200 Pro, TECAN) at 595 nm.

Protein (2 ml diluted to 4-5μg/ul in extraction buffer) was used for immunoprecipitation. 25 μl per sample of GFP-Trap (Chromotek) beads were equilibrated by washing 3 times with 1 ml of extraction buffer and then added to the protein sample. The sample was incubated for 1 hour at 4°C and then the beads were pulled down with a magnetic rack and supernatant was discarded. Beads were washed 3 times with extraction buffer and protein was eluted with 70 μl of 1x SDS-sample buffer pre-heated to 96°C, and then further incubated at 96°C for 5 minutes. Beads were removed and the sample was loaded onto SDS-PAGE and subject to western blot.

### Accession Numbers

Arabidopsis Genome Initiative locus identifiers for the genes mentioned in this article are as follows: HSFA1a (AT4G17750), HSFA1b (AT5G16820), HSFA1d (AT1G32330), HSFA1e (AT3G02990), WIND1 (AT1G78080), WIND2 (AT1G22190), ERF113_RAP2.6L (AT5G13330), ERF109 (AT4G34410), ERF115 (AT5G07310), SCL5 (AT1G50600), DOF3.4 (AT3G50410), PLT3 (AT5G10510), ZAT6 (AT5G04340), ZAT10_STZ (AT1G27730), ARR4 (AT1G10470), ARR6 (AT5G62920), ARR8 (AT2G41310), ARF19 (AT1G19220) and MP/ARF5 (AT1G19850).

## Acknowledgements

We thank Bart Rymen and Alice Lambolez for their technical help at an early stage of this research and Yu Chen for his helpful comments during the manuscript preparation. We also thank Ayami Furuta, Mariko Mouri, Chika Ikeda, Noriko Doi, and Akiko Hanada for their technical assistance.

## Author contributions

DC, AI and KS conceived the study and designed the experiments. DC and AK performed most of experiments with assistance from AI, AT, KEJ, MP, YK, DSF, TT, MI, TS, NO, and PAW. DC and KS wrote the draft of the manuscript. PAW, LDV and KYS reviewed and edited the manuscript. All authors discussed the results and commented on the manuscript.

## Supplementary Data

**Supplementary Figure S1.** Translocation of HSFA1d to nuclei after wounding.

**Supplementary Figure S2.** Evaluation of *hsfa1* mutants in wound-induced callus formation and *in vitro* shoot regeneration.

**Supplementary Figure S3.** Evaluation of *HSFA1dΔ1* overexpression or heat shock’s effect on shoot regeneration.

**Supplementary Figure S4.** Regulation of a large set of stress responsive genes by HSFA1

**Supplementary Figure S5.** Details of knockout and overexpression of HSFA1-target genes

**Supplementary Data Set 1.** A list of genes that are induced by wounding and heat stress.

**Supplementary Data Set 2.** A list of genes downregulated in *hsfa1abd-2* after wounding

**Supplementary Data Set 3.** A list of genes bound by HSFA1d in our ChIP-seq experiment.

**Supplementary Data Set 4.** A list of transcription factors regulated by HSFA1

**Supplementary Data Set 5.** A list of primers used in this study.

## Data Availability

The RNA-seq and ChIP-seq datasets produced in this study are available in the gene expression omnibus (GEO) under the series GSE306295 and GSE232157 respectively.

