## Supplemental Figures for "Wounding activates the HSFA1 transcription factors to promote cellular reprogramming in Arabidopsis"

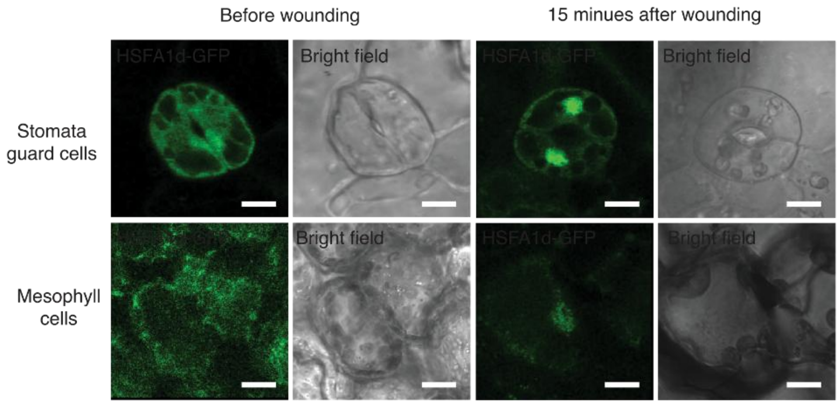


**Supplementary Figure S1. Translocation of HSFA1d to nuclei after wounding.** (Supports Figure 1)

Confocal images showing the subcellular localisation of HSFA1d-GFP proteins before and 15 minutes after wounding. Stomata guard cells and mesophyll cells from *pHSFA1d::HSFA1d-GFP hsfa1abd* cotyledons were visualized. Scale bars, 10 µm.

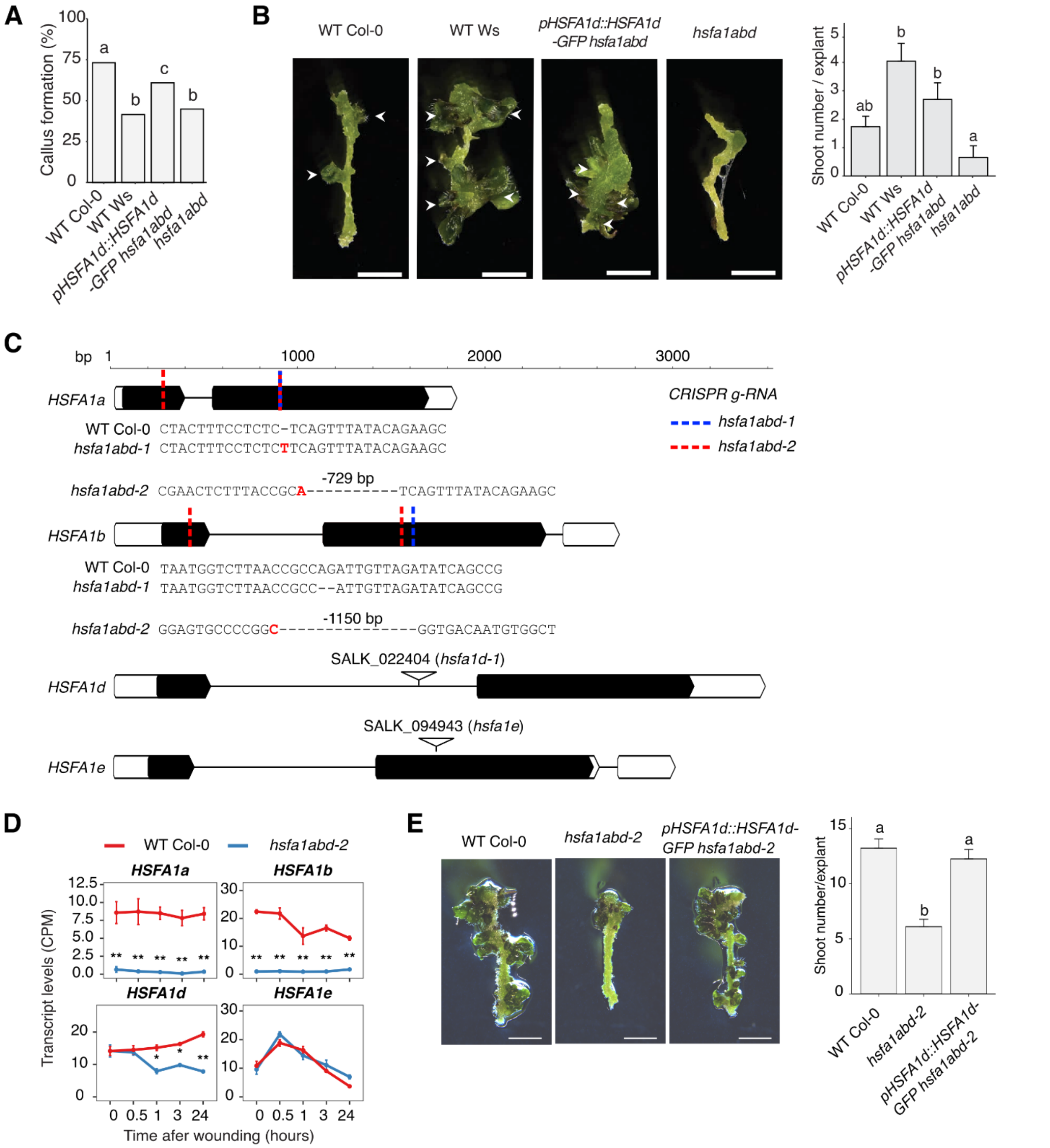


**Supplementary Figure S2. Evaluation of *hsfa1* mutants in wound-induced callus formation and *in vitro* shoot regeneration.** (Supports Figure 2)

(A) Quantification of the proportion of WT Col-0, WT Ws, *pHSFA1d::HSFA1d-GFP hsfa1abd* and *hsfa1ab* hypocotyls that form callus within one week after cutting (right). Sample sizes are WT Col-0 (*n* = 54), WT Ws (*n* = 60) *pHSFA1d::HSFA1d-GFP hsfa1bd* (*n* = 60), and *hsfa1abd* (*n* = 60). Different letters indicate significant differences based on pairwise two-proportions z-test ($P < 0.05$). (B) Images (left) and quantification (right) of shoot regeneration from hypocotyl explants of WT Col-0, WT Ws, *pHSFA1d::HSFA1d-GFP hsfa1abd* and *hsfa1abd*. All explants were incubated on CIM for 4 days and then on SIM for 14 days at 22°C. Arrows indicate regenerated shoots. Values represent mean number of shoots produced per explant, and error bars represent ± SE. Sample

sizes are WT Col-0 (*n* = 54), WT Ws (*n* = 60) *pHSFA1d::HSFA1d-GFP hsfa1bd* (*n* = 60), and *hsfa1abd* (*n* = 60). Different letters indicate significant differences based on one-way ANOVA and posthoc Tukey test ($P < 0.05$). Scale bars, 2 mm.
(C) Diagram showing the four HSFA1 member genes and the location of either gRNA target sequence or T-DNA location. Sequences of the edited loci in the CRISPR mutants of HSFA1a and HSFA1b are shown. Bases in red indicated insertions.
(D) Line plot showing the transcript levels of HSFA1 TF family members in WT and *hsfa1abd-2*. Data is from RNA-seq analysis. Error bars represent SE and asterisks indicate statistical differences based in Edge-R analysis (* $P < 0.05$, ** $P < 0.01$).
(E) Images (left) and quantification (right) of shoot regeneration from hypocotyl explants of WT Col-0, *hsfa1abd-2*, *pHSFA1d::HSFA1d-GFP* and *hsfa1abd-2*. All explants were incubated on CIM for 4 days and then on SIM for 14 days at 25°C. Arrows indicate regenerated shoots. Values represent mean number of shoots produced per explant, and error bars represent ± SE. Sample sizes are WT Col-0 (*n* = 33), *pHSFA1d::HSFA1d-GFP hsfa1abd-2* (*n* = 32), and *hsfa1abd-2* (*n* = 33). Different letters indicate significant differences based on one-way ANOVA and posthoc Tukey test ($P < 0.05$). Scale bars, 2 mm.

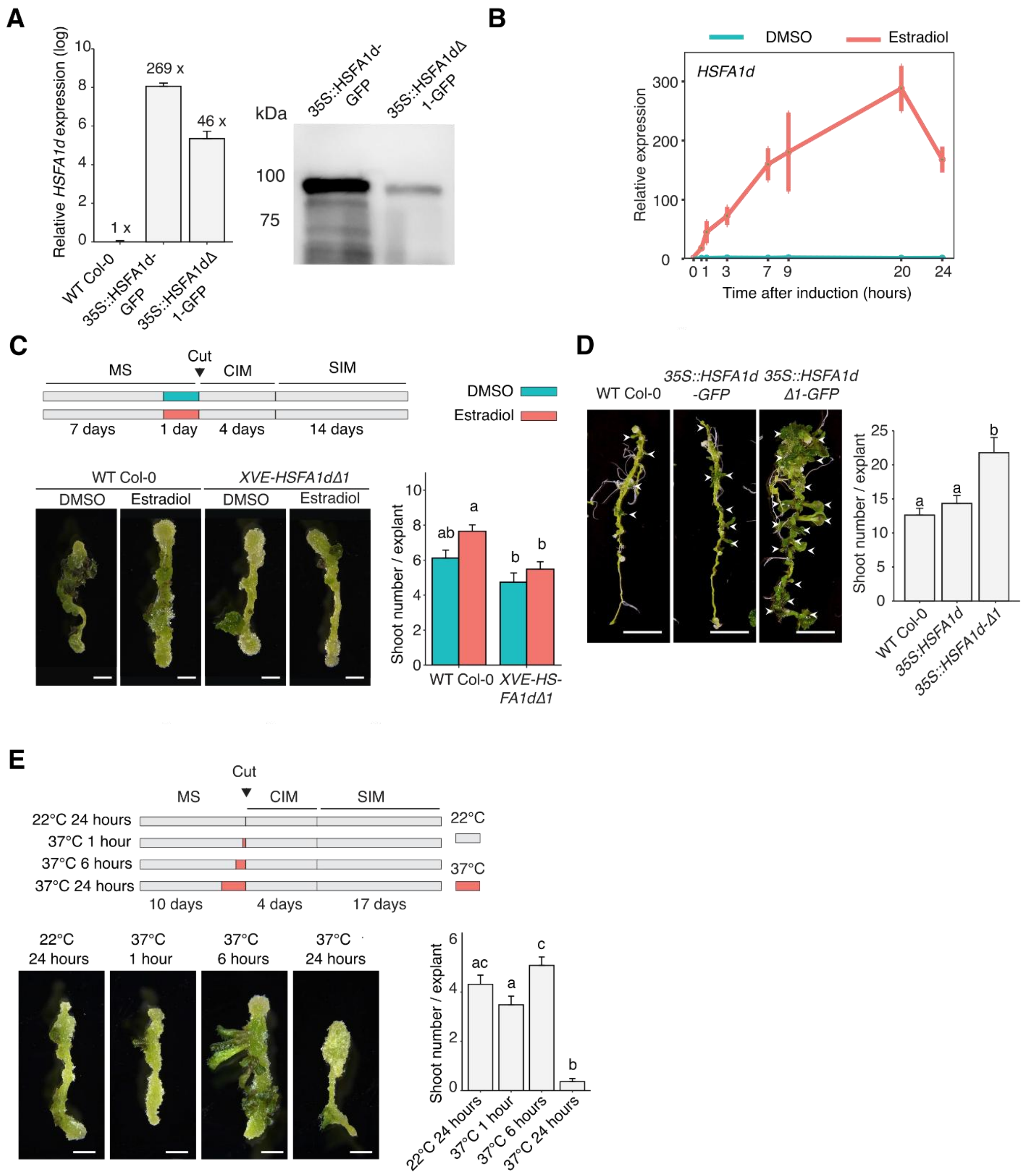


**Supplementary Figure S3. Evaluation of *HSFA1d*Δ*1* overexpression or heat shock's effect on shoot regeneration.** (Supports Figure 3)

(A) RT-qPCR analysis of *HSFA1d* transcripts (left) and western blot analysis of HSFA1 proteins (right) in *35S::HSFA1d-GFP* and *35S::HSFA1dΔ1-GFP* seedlings. RNA was extracted from several seedlings 17 days after germination grown on MS media. The expression levels were normalized to WT Col-0 level. Whole protein extracts from the overexpression lines were used and HSFA1d-GFP proteins were detected with anti-GFP antibodies. A total of 20 µg of proteins were loaded into each lane. Values represent log mean expression relative to PP2A3 normalised to WT Col-0 level, and error bars represent ± SE.

(B) RT-qPCR analysis showing the expression of *HSFA1d* upon addition of DMSO or 1 µM 17-ß-estradiol to *XVE-HSFA1dΔ1* seedlings. Values represent mean expression relative to PP2A3 normalised to time-0, and error bars represent ± SE.
(C) Images (left) and quantification (right) of shoot regeneration from root explants of WT Col-0, *35S::HSFA1d-GFP* and *35S::HSFA1dΔ1-GFP*. Sample sizes are WT Col-0 (*n* = 29), *35S::HSFA1d-GFP* (*n* = 27) and *35S::HSFA1dΔ1-GFP* (*n* = 28). Values represent mean number of shoots produced per explant, and error bars represent ± SE. Different letters indicate significant differences based on one-way ANOVA and posthoc Tukey test ($P < 0.05$). Scale bars, 2 mm.
(D) A diagram showing the experimental setup for the transient *HSFA1dΔ1* induction (top). Representative images (left) and quantification (right) of shoot regeneration from WT Col-0 and *XVE-HSFA1dΔ1* hypocotyl explants incubated in the presence of DMSO or 1 µM 17-ß-estradiol. Sample sizes are WT Col-0 treated with DMSO (*n* = 28), WT Col-0 treated with 17-ß-estradiol (*n* = 37), *XVE-HSFA1dΔ1* treated with DMSO (*n* = 36) and *XVE-HSFA1dΔ1* treated with 17-ß-estradiol (*n* = 25). Values represent mean number of shoots produced per explant, and error bars represent ± SE. Different letters indicate significant differences based on one-way ANOVA and posthoc Tukey test ($P < 0.05$). Scale bars, 2 mm.
(E) A diagram showing the experimental setup for the heat stress treatment (left). Representative images (middle) and quantification (right) of shoot regeneration from WT Col-0 hypocotyl explants. Sample sizes are 22°C (*n* = 59) and 37°C (*n* = 49-56). Values represent mean number of shoots produced per explant, and error bars represent ± SE. Different letters indicate significant differences based on one-way ANOVA and posthoc Tukey test ($P < 0.05$). Scale bars, 2 mm.

**A**

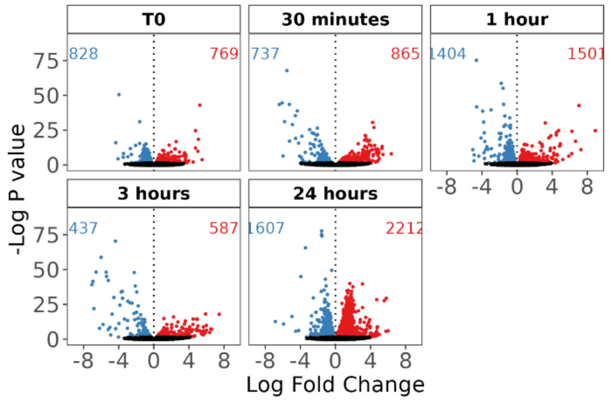


**B**

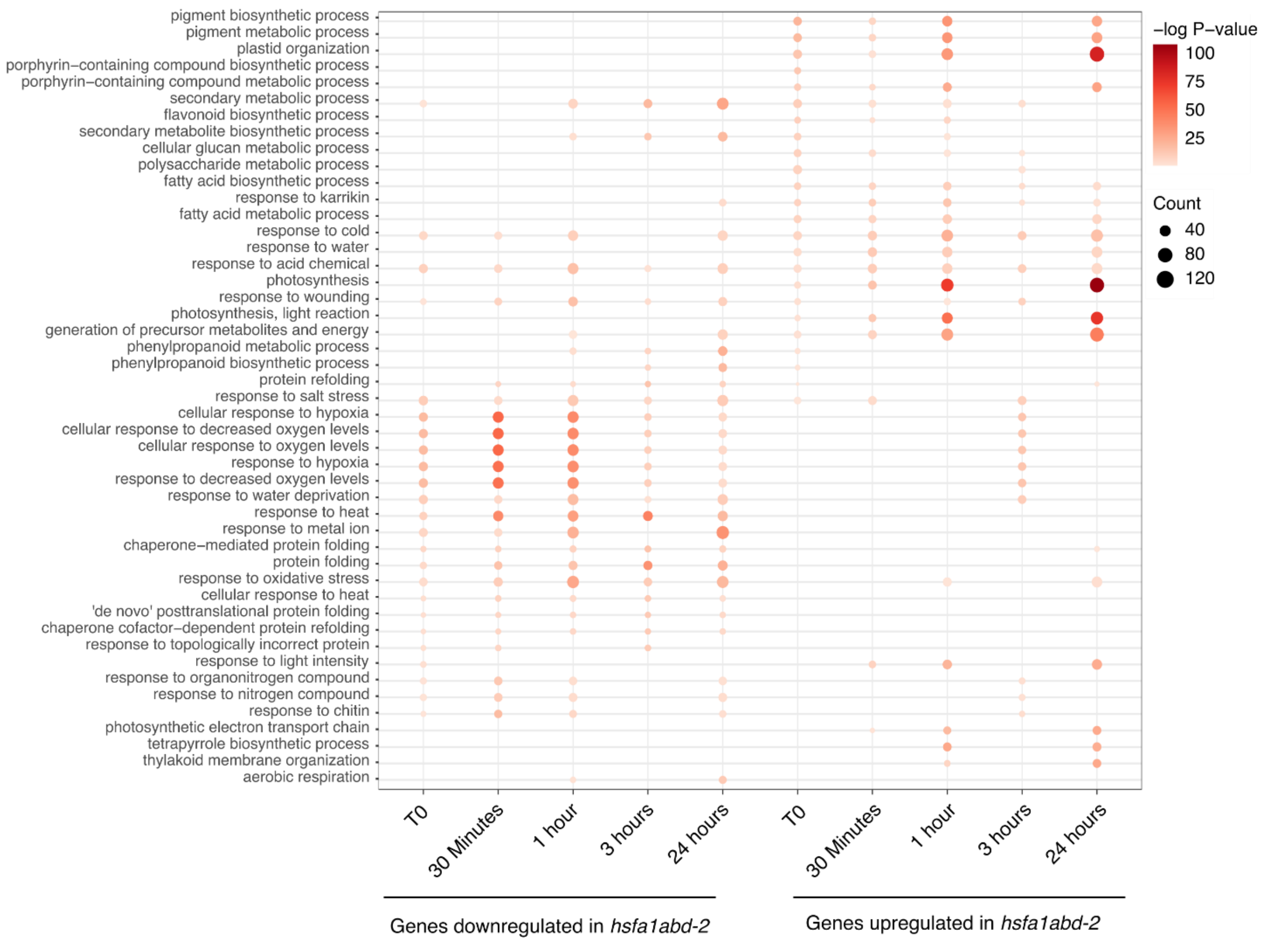


**Supplementary Figure S4. Regulation of a large set of stress responsive genes by HSFA1.** (Supports Figure 4)

(A) Volcano plot showing the differentially expressed genes (edgeR, false discovery rate [FDR] < 0.05) between WT Col-0 and *hsfa1abd-2* hypocotyls after wounding. Significantly upregulated or downregulated genes in *hsfa1abd-2* seedlings are depicted as red or blue dots, respectively. Red or blue numbers indicate the total number of upregulated or downregulated genes.

(B) GO enrichment analysis for genes differentially expressed in *hsfa1abd-2* seedlings. The most enriched GO categories among upregulated genes at any given time point.

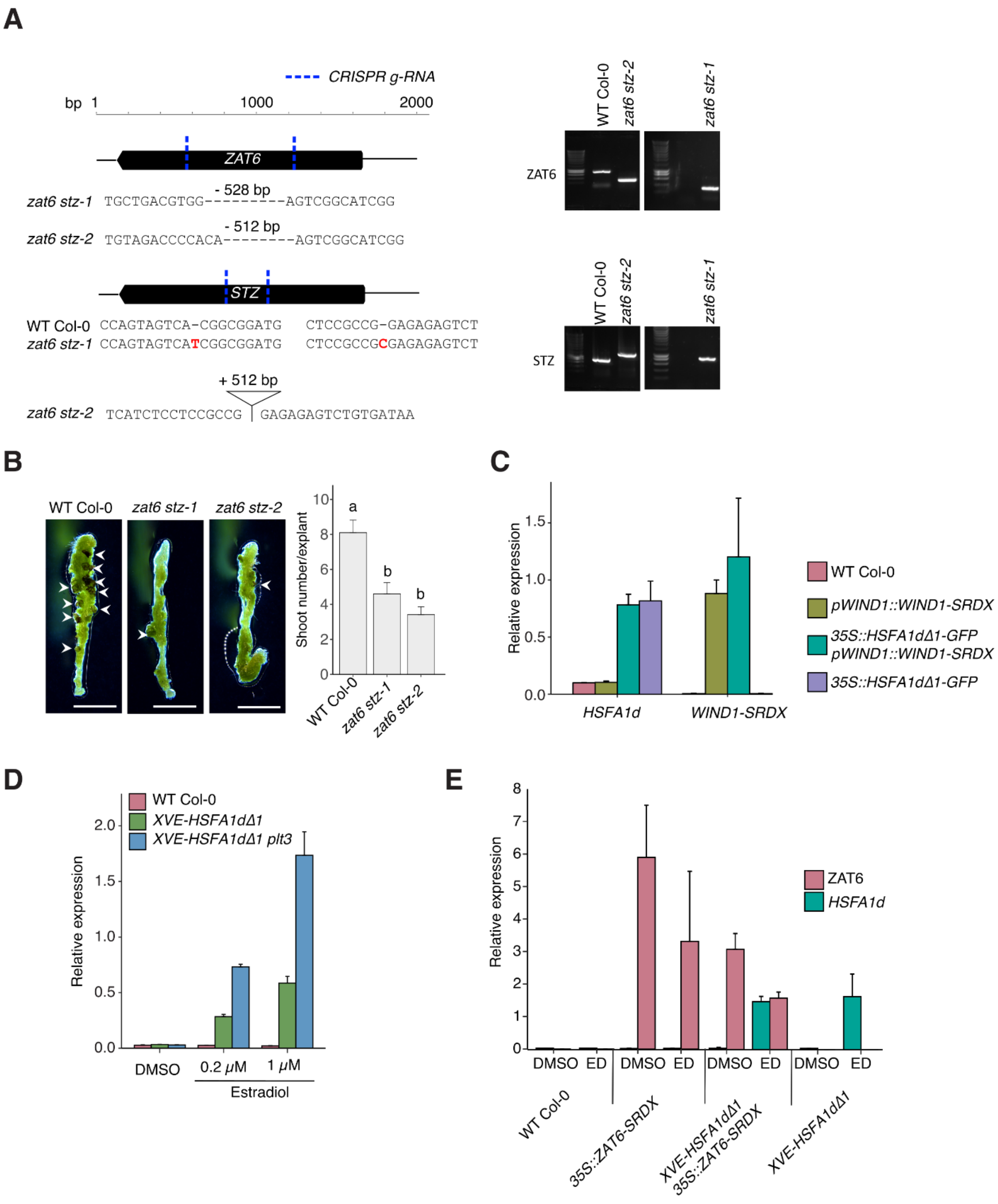


**Supplementary Figure S5. Details of knockout and overexpression of HSFA1 target genes.** (Supports Figures 4 and 6)

(A) Left; Diagram showing the location of target gRNAs (Supplementary Data Set 5) in the gene bodies of STZ and ZAT6 as well as the DNA sequence the mutated loci of mutants. All four gRNAs were cloned into a single vector *pFASTRK-pRPS5A-BP-AtCas9-BP*-G7T-A-BsaI-ccdB-BsaI-G* by golden-gate cloning as described (Develtere et al., 2024) and transformed into WT Col-0. T3 homozygous plants were used for analysis. Right; DNA electrophoresis gel showing PCR products from genomic-DNA of WT and mutant plants using primers listed in Supplementary Data Set 5.

Mutants used for phenotyping are labelled. Both double mutants have large deletions in *STZ* while zat6 *stz-1* has two 1bp insertions in *ZAT6* and a *zat6 stz-2* has a large insertion in *ZAT6*.
(B) Images (left) and quantification (right) of shoot regeneration from hypocotyl explants of WT Col-0, *zat6 stz-1* and *zat6 stz-2* at 25°C. Sample sizes are WT Col-0 ($n$ = 52), *zat6 stz-1* ($n$ = 48), and *zat6 stz-2* ($n$ = 48). Values represent mean number of shoots produced per explant, and error bars represent ± SE. Different letters indicate significant differences based on one-way ANOVA and posthoc Tukey test ($P < 0.05$). Scale bars, 2 mm.
(C) RT-qPCR analysis of *HSFA1d* and *WIND1-SRDX* expression levels in WT-Col-0, *XVE-HSFA1dΔ1*, *pWIND1::WIND1-SRDX*, and *XVE-HSFA1dΔ1 pWIND1::WIND1-SRDX* plants. RNA was extracted from several leaves of 2-week-old seedlings grown on 1 µM 17-ß-estradiol. Expression of each gene is relative to the *PP2A3* housekeeping gene. Values represent mean expression relative to *PP2A3* and error bars represent ± SE.
(D) RT-qPCR analysis of HSFA1d in WT Col-0, *XVE-HSFA1dΔ1* and *XVE-HSFA1dΔ1 plt3* plants. RNA was extracted from 2-week-old seedlings grown on MS containing either DMSO, 0.2 µM or 1 µM 17-ß-estradiol. Expression levels are relative to *PP2A3* and error bars represent ± SE.
(E) RT-qPCR analysis of *HSFA1d* and *ZAT6-SRDX* in WT-Col-0, *XVE-HSFA1dΔ1*, *35S::ZAT6-SRDX*, and *XVE-HSFA1dΔ1 35S::ZAT6-SRDX* plants. RNA was extracted from whole 2-week-old seedlings treated with DMSO or 10 µM 17-ß-estradiol. Expression levels are relative to *PP2A3* and error bars represent ± SE.
